# Dynamic changes in chromosome and nuclear architecture during maturation of normal and ALS C9orf72 motor neurons

**DOI:** 10.1101/2025.09.22.677835

**Authors:** Özgün Uyan, Snehal Sambare, Marlies E. Oomen, Nicholas Wightman, Allana Schooley, Joseph R. Klim, Houda Belaghzal, Özkan Aydemir, Betul Akgol-Oksuz, Zeynep Sena Agim Uslu, Kevin Eggan, Robert H. Brown, Job Dekker

**Affiliations:** Department of Neurology, University of Massachusetts Chan Medical School, Worcester, MA 01605, USA; Department of Systems Biology, University of Massachusetts Chan Medical School, Worcester, MA 01605-0103, USA; Department of Stem Cell and Regenerative Biology, Harvard University, Cambridge, MA 02138, USA; Stanley Centre for Psychiatric Research, Broad Institute of MIT and Harvard, Cambridge, MA 02142, USA; Program in Molecular Medicine, University of Massachusetts Chan Medical School, Worcester, MA 01605, USA; Howard Hughes Medical Institute, Chevy Chase, MD, USA

## Abstract

We have investigated changes in chromosome conformation, nuclear organization, and transcription during differentiation and maturation of control and mutant motor neurons harboring hexanucleotide expansions in the *C9orf72* gene that cause amyotrophic lateral sclerosis (ALS). Using an *in vitro* reprogramming, differentiation and neural maturation protocol, we obtained highly purified populations of post-mitotic motor neurons for both normal and diseased cells. As expected, as fibroblasts are reprogrammed into iPSCs, and as iPSCs differentiate into motor neurons, chromatin accessibility, chromosome conformation, and nuclear organization change along with large-scale alterations in transcriptional profiles. We find that the transcriptome changes extensively during the first three weeks of post-mitotic neuronal maturation, with thousands of genes changing expression, but then is relatively stable for the next three weeks. In contrast, chromosome conformation and nuclear organization continue to change over the entire 6-week maturation period: chromosome territoriality increases, long-range interactions along chromosomes decrease, compartmentalization strength increases, and centromeres and telomeres increasingly cluster. In motor neurons derived from ALS patients such changes in chromosome conformation were much reduced. Chromatin accessibility changes also showed delayed maturation. The transcriptome in these cells matured relatively normally but with notable changes in expression of genes involved in lipid, sterol and mitochondrial function. We conclude that neural maturation is associated with large scale post-mitotic changes in gene expression, chromosome conformation and nuclear organization, and that these processes are defective in motor neurons derived from ALS patients carrying *C9orf72* hexanucleotide repeat expansions.

## Introduction

Amyotrophic lateral sclerosis (ALS) is a fatal motor neuron disease that is caused by the degeneration of upper and lower motor neurons located in the brain and spinal cord (Brown and Al-Chalabi, 2017; Rowland and Shneider, 2001). In typical cases of ALS, symptoms start with adult-onset focal muscle weakness of limb muscles leading to widespread paralysis; death occurs within 3-5 years due to respiratory failure after the first symptoms are observed. Although ALS has classically been defined as a pure motor neuron disease, it is now recognized as a multigenic neurodegenerative disorder in which both motor neurons and other cell types contribute to motor neuron death (Haidet-Phillips et al., 2011; Haukedal and Freude, 2019; Smethurst et al., 2020).

Recent advances in genetic technologies have revealed multiple genes that contribute to the initiation and progression of the disease (Udine et al., 2023; Wang et al., 2023). Discovery of these genes (e.g., *SOD1*, *FUS*, *TARDBP* (TDP-43), *C9orf72*) has enhanced our knowledge of biological pathways that are potentially affected in ALS.

The most common cause of ALS is the hexanucleotide repeat (GGGGCC; G_4_C_2_) expansion (HRE) in the first intron of the chromosome 9 open reading frame 72 (*C9orf72*) gene. This mutation accounts for ∼40-50% of familial ALS (fALS) and ∼10% of sporadic ALS (sALS) cases. In healthy individuals, the number of G_4_C_2_ hexanucleotide repeats varies from 2 to roughly 25 or 30. However, in C9orf72 cases (ALS or frontotemporal dementia (FTD)) this is expanded to hundreds or even thousands of repeats; the repeat size can vary between ∼500 bp and ∼25 kbp in length (DeJesus-Hernandez et al., 2011; Renton et al., 2011). Three possible mechanisms have been proposed for how the HRE in *C9orf72* contributes to ALS pathology. (1) Bidirectional transcription of the *C9orf72* gene within the promoter and intron 1 has been shown to generate G_4_C_2_ sense and G_2_C_4_ antisense expanded RNAs (Zu et al., 2013). These intronic transcripts including the HRE can give rise to G-quadruplex and hairpin structures that form RNA foci in the nucleus (several to many) or cytoplasm (rare) and sequester RNA-binding proteins (Conlon et al., 2016; Haeusler et al., 2014; Ramesh et al., 2020). (2) HRE containing RNAs can be translated through repeat-associated, non-ATG (RAN) translation, resulting in the synthesis of five different dipeptide protein repeats (DPRs) that may aggregate in the cytoplasm and nucleus (Lagier-Tourenne et al., 2013; Mori et al., 2013; Zu et al., 2013), and (3) the presence of the HRE can cause abortive and aberrant transcription of the C9orf72 gene itself, leading to decreased expressions of the *C9orf72* gene at the RNA and protein levels (DeJesus-Hernandez et al., 2011; Gijselinck et al., 2012; Saberi et al., 2018; Xiao et al., 2015; Zhu et al., 2020). However, even after a decade of study, there is still not a consistent view of which pathways are most important in initiating and sustaining C9orf72-related motor neuron death. Recent results also suggest that both loss-of-function and gain-of-function mechanisms may act synergistically to initiate disease pathology (Boivin et al., 2020; Lutz, 2020; Shao et al., 2019; Shi et al., 2018; Zhu et al., 2020).

Advancements in human induced pluripotent stem cell (hiPSC) research (Takahashi and Yamanaka, 2006) have facilitated the modeling of unique cell types *in vitro*, producing homogeneous populations of specific cell types. Over the years, numerous motor neuron differentiation protocols have been developed and optimized (Devlin et al., 2015; Dimos et al., 2008; Kiskinis et al., 2014; Klim et al., 2019; Maury et al., 2015). Depending on the protocol, maturation time for neurons varies from days to months (Klim et al., 2019; Lopez-Gonzalez et al., 2019). Diverse aspects of motor neuron biology can be assessed using these populations of differentiated motor neurons, some reflecting characteristics of motor neurons *in situ*. Using longer maturation periods allows better assessment of various physiological properties such as viability, action potential, and synaptic input (Devlin et al., 2015).

In addition to the physiological changes that occur during cellular differentiation, several molecular changes also occur. One such change is in the 3D organization of the genome, which facilitates gene expression changes necessary for cell state switches. The 3D organization of chromosomes, and changes in that organization, can be assessed by Hi-C methods (see (Jerkovic and Cavalli, 2021) and (Dekker and Mirny, 2024) for recent reviews). At the scale of the nucleus, chromosomes are found to occupy distinct territories, with some overlap at their peripheries (Branco and Pombo, 2006) but with limited topological intermingling (Tavares-Cadete et al., 2020). Chromosomes are compartmentalized at the scale of hundreds of kilobases (kb) up to megabases (Mb), where chromatin domains of different types (e.g., eu-and heterochromatin, or A-and B compartments (Lieberman-Aiden et al., 2009; Rao et al., 2014; Spracklin et al., 2023)) form preferentially homotypic interactions so that chromatin domains of different types spatially segregate. At a smaller scale, tens to hundreds of kb, chromatin loops form through affinity driven interactions, e.g., between promoters and other active elements (Goel et al., 2023; Schooley et al., 2024; Zhang et al., 2019a), and through active loop extrusion, e.g., by cohesin leading to dynamic loops throughout the genome, and positioned loops at sites where cohesin stalls such as CTCF-bound sites (Fudenberg et al., 2017). Topologically Associating Domains (TADs) (Dixon et al., 2012; Nora et al., 2012) emerge as a population average feature driven by cohesin-mediated loop extrusion with TAD boundaries reflecting positions where cohesin is stalled (Rao et al., 2017).

The 3D organization of chromosomes changes as cells differentiate, in a manner that is correlated with changes in gene expression. For example, during differentiation of embryonic stem cells (ESCs) into neural progenitor cells (NPCs) and terminally differentiated neurons, there are dynamic changes in 3D positioning of genes leading to changes in the amount of transcriptionally active/open (A) compartment versus inactive/closed (B) compartment states (Bonev et al., 2017; Phillips-Cremins et al., 2013; Rajarajan et al., 2018). ESCs have globally active chromatin states, while more differentiated cells such as NPCs and post-mitotic neurons demonstrate diverse condensed chromatin structures depending on the cell type, differentiation, and maturation states (Hawkins et al., 2010; Kishi et al., 2012; Nothof et al., 2022). These types of changes can also be seen using transposase-accessible chromatin with high-throughput sequencing (ATAC-seq), and DNase I hypersensitivity sequencing (DNase-seq), which also show dynamic changes in chromatin accessibility during neural differentiation (de la Torre-Ubieta et al., 2018; Frank et al., 2015; Melendez-Ramirez et al., 2021). These dynamic changes have a crucial role during brain development (Trevino et al., 2020). Studies using Hi-C showed that spatial chromatin interactions are reorganized during human brain development using cells from different regions of the central nervous system, including neural progenitors and post-mitotic neurons (Heffel et al., 2024; Won et al., 2016). However, despite these advances, it remains unknown how these critical dynamic events in 3D arrangements of chromatin and chromosomes may be perturbed in neurodegenerative disorders.

In this study, we used Hi-C, RNA-seq, and ATAC-seq to investigate the relation between spatial genome organization and the transcriptome during motor neuron differentiation and maturation in healthy and disease (ALS) conditions. We identified global re-organization of higher order chromatin structures such as A/B compartment switches and changes in compartment strengths. While we detected a decrease in the frequency of total long-range compartmental interactions in mature motor neurons, we find increased preference for compartmental homotypic interactions, both in cis and in trans. Further, we observed progressive increases in enriched telomere-telomere and centromere-centromere long-range cis-and trans-chromosomal interactions as motor neurons mature, suggesting large-scale post-mitotic re-organization of chromosome territories and nuclear organization. Comparing healthy control cells to ALS patient-derived cells with the *C9orf72* HRE mutations, we find that compared to control, the normal transcriptional state is only partically established in ALS neurons and that the nuclear organization of fully matured differentiated motor neurons is not achieved, suggesting that genome organization in ALS-derived motor neurons, at least when differentiated *in vitro*, remains in an immature state.

## Results

### Generating motor neurons from primary fibroblasts

To generate induced pluripotent stem cells (iPSC)-derived motor neurons, primary healthy dermal fibroblasts (PF) were reprogrammed into iPSCs using an integration-free reprogramming kit (Figure S1A and B; see methods). After validating iPSC properties by immunofluorescence (Figure S1C), iPSCs were then differentiated into motor neurons (MN) using a fluorescence-activated cell sorting (FACS)-based differentiation method to obtain fully differentiated, homogeneous post-mitotic immature neurons (Klim et al., 2019) (Figure 1A and 1B). FACS-based selection of motor neurons allowed us to eliminate different cell types such as glial and NPCs that are unintentionally generated during the differentiation process. Motor neuron maturation was then induced for 6 weeks by addition of three neurotrophic factors: brain-derived neurotrophic factor (BDNF), glial-derived neurotrophic factor (GDNF), and ciliary neurotrophic factor (CNTF) (Figure 1B).

**Figure 1.**
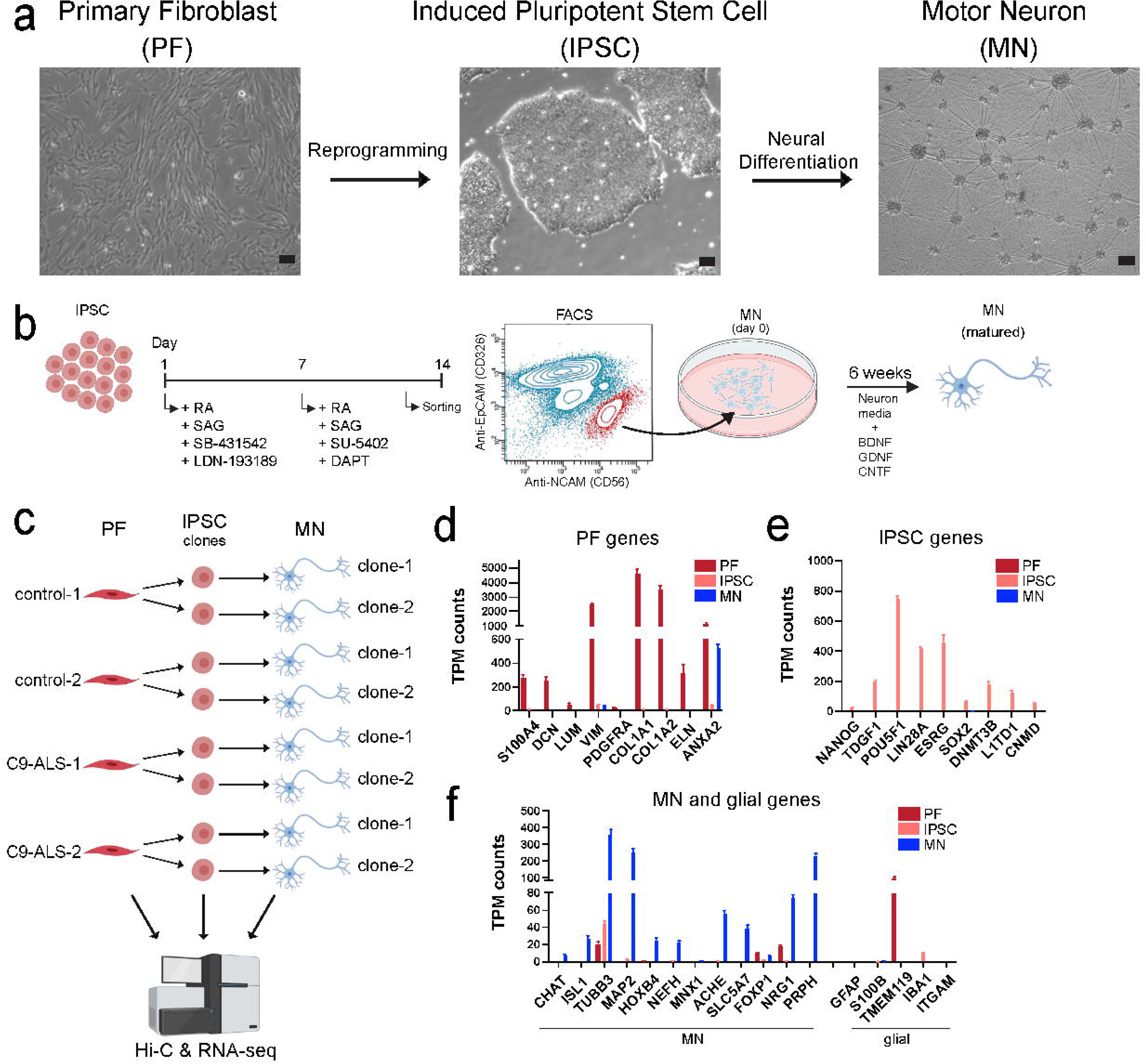
Generating motor neurons from primary fibroblasts via reprogramming and neural differentiation/maturation. a) Brightfield images of three different cell-types (scale bar=100 μm). b) Schematic view of neural differentiation from IPSCs into motor neuron and motor neuron maturation. IPSCs are treated with listed small molecules for 14 days. At day 14, cells were dissociated and sorted. NCAM+ and EPCAM-cells (population highlighted in red) were collected and plated as immature post-mitotic neurons that were cultured 6 more weeks for maturation (SB-431542: TGF-beta/Smad inhibitor; LDN-193189: BMP pathway inhibitor; RA: retinoic acid; SAG: Smoothened agonist; DAPT: γ-secretase inhibitor; SU-5402: FGFR1 inhibitor; BDNF: brain-derived neurotrophic factor; GDNF: glial-derived neurotrophic factor; CNTF: ciliary neurotrophic factor. c) Schematic view of the study design: primary fibroblasts of two healthy controls and two ALS patients with C9orf72 mutation were reprogrammed into IPSCs. Two independent IPSC clones for each line were picked for neural differentiations. Hi-C and RNA-seq libraries were prepared for each clone. d) TPM counts of PF related genes were shown for PF (red), IPSC (salmon) and MN (blue) cell-types. e) TPM counts of IPSC related genes were shown for PF, IPSC and MN cell-types. f) TPM counts of MN and glial related genes were shown for PF, IPSC and MN cell-types.

We used the above differentiation protocol to generate iPSCs from two independent control fibroblast cell lines (from two normal donors). We then chose two clones from each iPSC population to generate MNs (Figure 1C). To validate that each population was generated as intended, we analyzed the transcriptomic profiles for all generated lines (Figure 1D-F). Focusing on a selected set of cell type-specific genes, we find that these showed expression only in the expected cell types, confirming that pure populations of motor neurons were generated that were not contaminated by other cell-types such as glial cells during differentiation or after sorting (Figure 1F).

### Dynamics of chromosome compartmentalization during reprogramming, neural differentiation, and maturation

We used Hi-C to examine genome-wide compartmentalization in PFs, iPSCs and differentiated MNs. In chromosome-wide Hi-C interaction maps, compartmentalization of chromosomes in A and B compartments is reflected in a “checkerboard” pattern of enriched and depleted interactions. Visual inspection of Hi-C interaction maps shows obvious changes in compartmentalization patterns between PFs, iPSCs and MNs (Figure 2A). First, the checkerboard pattern is sharp and obvious in PFs, while in iPSCs and MNs such patterns, while clearly present, are composed of less frequent long-range interactions. Second, the precise pattern of the compartmental interactions along chromosomes differs in the different cell types, indicating numerous loci are in different compartment types in these different cell types.

**Figure 2.**
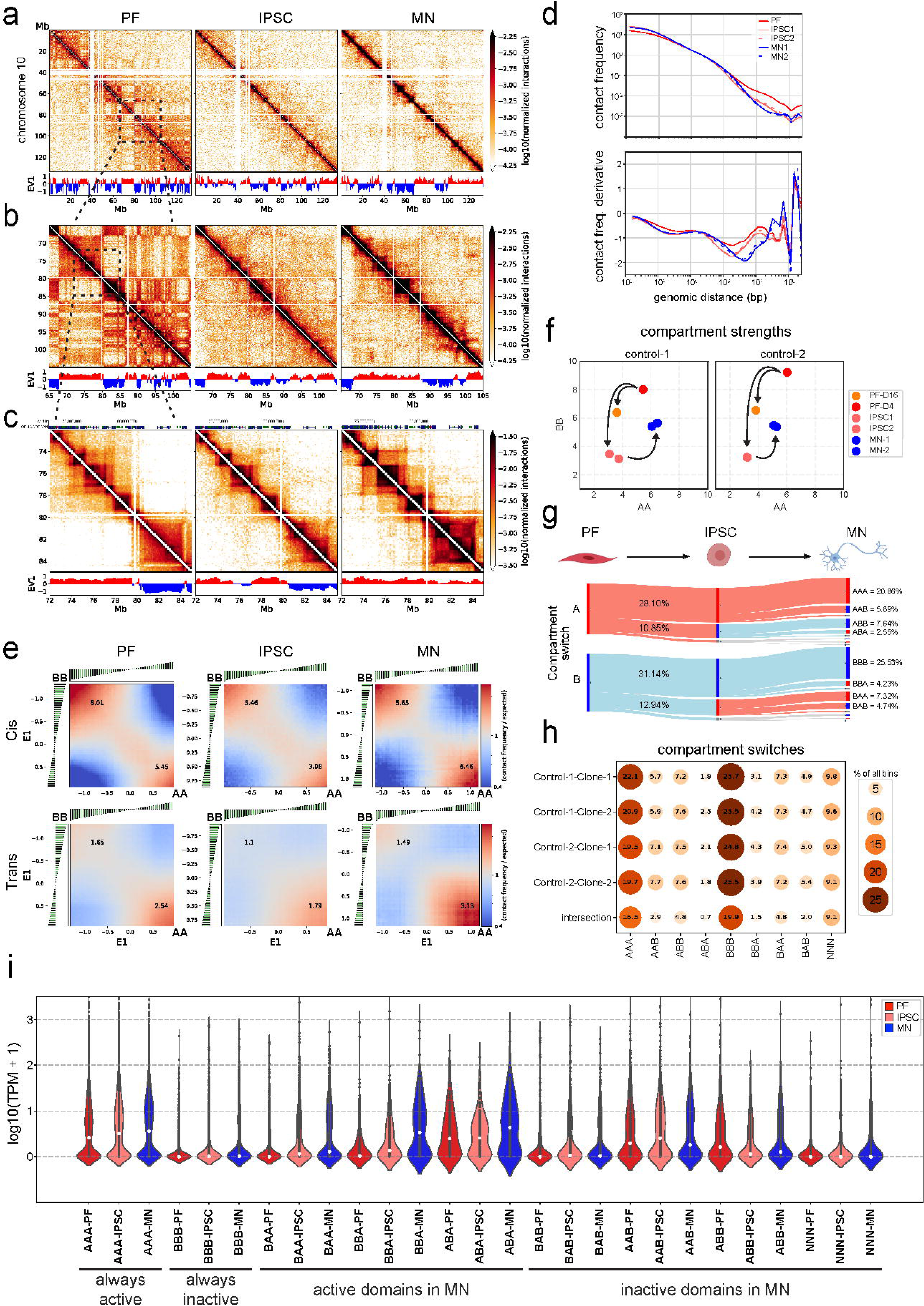
Reorganization of chromosome conformation during reprogramming, neural differentiation and maturation. a. Hi-C contact matrices of chromosome 10 at 100 kb resolution with A-B compartment tracks of first eigen value (EV1) in PFs, IPSCs and MNs. b. Zoomed in Hi-C region of chromosome 10 between 65-105 Mb at 100 kb resolution with A-B compartment tracks (EV1). c. Example Hi-C region (chromosome 10: 72-85 Mb) for TAD and looping structures with A-B compartment tracks in PFs, IPSCs and MNs. Top panel shows gencode genes (dense) for the region. d. Genome-wide contact frequency as a function of genomic distance between interaction pairs for PF (red), IPSCs (salmon) and MNs (blue) (1kb-∞) (top panel). Derivative plot of the contact frequency (bottom panel). e. Cis and trans saddle plots of genome-wide interactions for PFs, IPSCs and MNs. f. Compartment strengths of AA and BB were quantified using cis interactions obtained from saddle plots of PFs, IPSCs and MNs from control individuals. g. Eight major groups were detected in the analysis of compartment switches observed among PF, IPSC and MN during reprogramming, neural differentiation and maturation for clone #2 of control #1. First letter stands for PF, second is for IPSC and third is for MN. A is for euchromatin (EV1>0.05) and B is for heterochromatin (EV1<-0.05). h. Summary of all compartment switches observed in four control clones of PF, IPSC and MN. i. Violin plots demonstrate the TPM values of PF, IPSC and MN lines obtained from RNA-seq data for genes positioned within each category in figure 2h.

We quantified compartmentalization in different ways. First, the checkerboard pattern is routinely quantified using principal component analysis of the Hi-C interaction matrix, and the first Eigenvector (EV1) typically represents the compartment pattern (Lieberman-Aiden et al., 2009). EV1 shows clear differences between the three cell types (Figure 2A-C). Second, to quantify the frequency of compartmental interactions, we plotted interaction frequency *P* as a function of genomic distance *s* (Figure 2D). *P*(*s*) curves show the expected inverse correlation between interaction frequency and genomic distance between loci. Importantly, these plots show a clear reduction of interaction frequencies for loci separated by more than 1-2 Mb in iPSCs and MNs as compared to PFs (right of the dotted lines). This is the distance scale where interactions between compartment domains dominate. Third, we calculated the strength of compartmentalization, i.e., the strength of the preference of A domains to interact with A domains, and B domains with B domains. This is routinely calculated by “saddle plot” analysis (Nora et al., 2017). Saddle plots are Hi-C interactions maps where loci along both axes are sorted by their first eigenvector (EV1). Compartmentalization strength is then calculated by dividing the average contact enrichment between B-B and A-A pairs by the average A-B contact enrichment of the top 20% sorted eigenvectors (Nora et al., 2017). Compartmentalization strength quantifies the preference of formation of homotypic (A-A, and B-B) interactions as compared to heterotypic (A-B) interactions. We find that during reprogramming from PF to iPSC the strength of intra-chromosomal (cis) compartmentalization decreased (Figure 2E and 2F).

However, as iPSCs differentiate to MNs, this compartmentalization increases again for both A-A and B-B interactions. A-A interaction strength in MNs is comparable to that in PFs, while B-B interaction strength is somewhat lower (Figure 2F). Inter-chromosomal (trans) compartmentalization showed similar dynamics (Figure 2E). Thus, while the absolute number of compartmental interactions decreases as iPSCs differentiate and mature into MNs, the preference for homotypic A-A and B-B interactions becomes stronger.

Compartmentalization strength has been shown to be strongly reduced during mitosis (Naumova et al., 2013) and then is only slowly reestablished in the subsequent G1 phase (Abramo et al., 2019). Therefore, we tested the possibility that the increased compartmentalization strength in MNs as compared iPSCs was simply due to the fact that MNs had been post-mitotic for several weeks. To this end, we maintained fibroblasts for an additional period of 12 days in culture and performed Hi-C on PFs on day 4 (PF-D4) and day 16 (PF-D16). During this time fibroblasts became confluent and stopped dividing. Longer incubation of PFs led to a decrease in B-B and A-A compartmentalization strength on day 16 (Figure 2F).

Therefore, since longer incubation of post-mitotic PFs resulted in a reduction of compartment strength, the prolonged incubation period during MN maturation was probably not a factor in the observed increased compartmentalization strength in MNs.

Finally, we determined how A and B compartmentalization patterns along the genome changed as cells were reprogrammed and then differentiated into motor neurons. Overall, we detected 8 major compartment switching groups in PF, IPSC and MN lines of each clone (Figure 2G). For instance, the AAA group refers to genomic loci (100 kb bins) that are always compartment type A (active and open euchromatin) in PFs, iPSCs, and MNs. This category represents 16.5% of the genome. Likewise, 19.9% is always compartment type B (“BBB”, closed and inactive heterochromatin). Then, we compared all clonal control lines, and we found that 62.2% (the sum of all % of bins in the category “intersection”, Figure 2H) of all genomic bins showed the same type of grouping among PF, IPSC and MN lines in different clones of two control lines. Nearly 1/3 of all detectable bins of the genome do not change their compartment status (AAA or BBB) due to cell-type or between cell lines/clones (Figure 2H). Conversely, ∼16% of detectable bins show consistent dynamic compartment switches among individuals and cell-types. The remaining genomic bins show some variation between clones and individuals. All possible switching orders were observed (e.g. ABA, AAB, BBA, etc.) with comparable frequencies (∼6-9% of bins for each order, Figure 2H). These results indicate prominent dynamics in compartmentalization during differentiation and maturation in addition to some variation due to differences between donors and derived clonal lines.

To determine how transcriptional changes are related to compartment switches, we analyzed expression levels of genes (TPM: transcripts per million reads) that reside within the intersecting compartments (loci that show consistent compartmentalization between individuals and clones) for each switch group for each of the three cell lines. We find that changes in expression levels are generally consistent with switches in compartment status. Genes in regions that are consistently in A domains are expressed at higher levels in all three cell types, while genes consistently located in B domains are expression at the lowest level, for each cell type. Expression of genes located in regions that consistently switch compartment types across the three cell types typically changes accordingly. Further, there are global gene expression level differences such that genes generally show higher expression pattern in MNs compared to PFs and IPSCs. One reason for this is that normalization by total reads is sensitive to the variance of gene expressions (Zhao et al., 2020).

To further support these findings, we analyzed chromatin architecture associated with cell type-specific gene expression. Using regions of 4 Mb surrounding genes of interest, we compared Hi-C maps for three PF, three iPSC, six MN, and four glial associated gene regions in the various cell types. Some gene regions showed substantial reorganization of compartments, TAD structures and looping interactions between PFs, iPSC and MN lines (Figure S2A and S2B) which correlate with gene expression profiles. For instance, the *COL1A2* gene is highly expressed and resides in a euchromatic A compartment domain in PFs. However, the gene shows no expression in iPSCs and MNs and is then located in heterochromatic B compartment domains. Similarly, the *MAP2* gene which is highly expressed in MNs but not in PFs and iPSCs, is located in a euchromatic A compartment domain in MNs but in a heterochromatic B compartment domain in PFs and iPSCs. Likewise, glial genes which are mostly not expressed in any of the three cell-types studied here, do not show major differences in genome architecture among these cell-types (Figure S2D). For some other genes such as *COL1A1*, *L1TD1*, *NEFH* and *PRPH* that show cell-type specific expression, we could not detect major differences in chromatin structure or chromatin state (A or B compartments) between the cell-types (Figure S2C). These genes reside within highly gene rich regions and therefore it is possible that expression of these other gene/gene clusters might be driving the chromatin structure and compartment status at the locus. Along these same lines, genes that are located in gene poor regions, could be the driving force for 3D architecture within these regions contributing to the cell-type specificity in PFs, iPSCs and MNs (Figure S2A and S2B).

### Motor neurons display prominent intra-and inter-chromosomal telomeric, centromeric interactions

We examined how inter-chromosomal interaction patterns change as cells are reprogrammed from fibroblasts into iPSCs, and during subsequent differentiation into mature motor neurons. First, visual inspection of Hi-C maps shows that although inter-chromosomal interactions are readily detectable in PF, iPSC, and MN cells (Figure 3A), MNs have the lowest level. This is visible by the depletion of interactions in the interchromosomal sections of the Hi-C maps of MNs as compared to the other cell types. This was confirmed by quantifying the fraction of intra-chromosomal (cis) genome-wide interactions (Figure 3B): the percentage of cis interactions was around 70% in PFs and iPSCs, while it increased to ∼85% in MN cells. This suggests that chromosome territoriality becomes more pronounced in MN cells.

**Figure 3.**
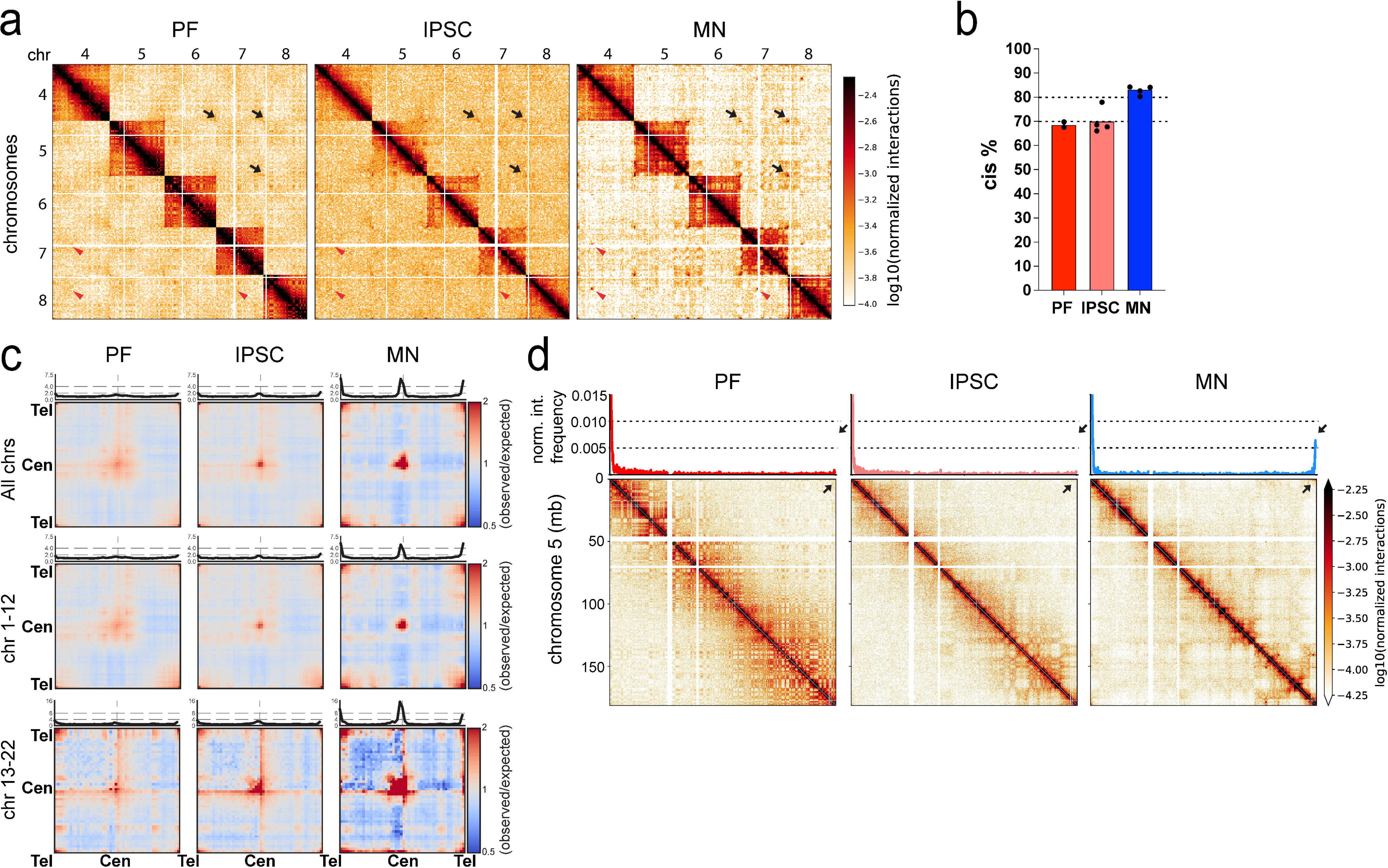
Motor neurons display prominent intra-and inter-chromosomal telomeric and centromeric interactions. a. Hi-C matrices show the inter-chromosomal (trans) interactions among chromosomes 4-8 in PF, IPSC and MN. Arrows show telomeric and centromeric interactions among PF, IPSC and MN cell-types (black arrows: telomere-telomere interactions, red arrowheads: centromere-centromere interactions) (resolution= 1Mb). b. Genome-wide intra-chromosomal (cis) interaction percentages in PF, IPSC and MN samples. c. Saddle plots of aggregated genome-wide trans interactions of telomere-telomere, centromere-centromere and telomere-centromere for all, large and small chromosomes of PF, IPSC and MN. Top panel shows the diagonal interaction quantifications. d. Hi-C matrices show cis interactions in chromosome 5 among PF, IPSC and MN cell-types (100kb bin size). Top panel demonstrates 4C interaction plots of the p-arm telomeric region throughout chromosome 5. Black arrows indicate cis interactions of p-and q-arm telomeric regions.

Second, we noticed in motor neuron Hi-C maps that despite a reduction in total inter-chromosomal interactions in general, a much pronounced “checkerboard” of compartmental interactions was present. Saddle plot analysis confirmed that inter-chromosomal (trans) A-A interactions became much more prominent in MNs as compared to PFs and iPSCs (Figure 2E). The dynamics of compartmentalization in trans mirrored that what was observed for cis compartmentalization where compartmentalization becomes weaker during reprogramming PFs into iPSCs, and then becomes much stronger, especially the preference of A-compartment domains to homotypically interact (above).

Third, we observed prominent interactions between telomeres and between centromeres of different chromosomes (black arrows and red arrows, respectively in Figure 3A). To quantify this, we calculated the average, and scaled to a universal size, inter-chromosomal interaction maps (Figure 3C). We observed clear preferential interactions between centromeres in all three cell types, but these became stronger in motor neurons, especially among the shorter chromosomes (Figure 3C, bottom row). To visualize this in another way, we plotted the interactions along the diagonal of these scaled inter-chromosomal interaction maps (shown in top panels of the interaction maps in Figure 3C), which reveals an obvious peak at the centromere-centromere positions. This peak was small in PFs and iPSCs but was much stronger in MNs.

Interestingly, these line plots also display peaks at the telomere-telomere interactions. These inter-chromosomal telomeric interactions also increased in frequency in MNs. We did not notice enriched centromere-telomere interactions.

Interestingly, interactions between telomeres of the same chromosome also increased in frequency in MNs as compared to PFs and iPSCs (Figure 3D). To explore such intra-chromosomal telomere-telomere interactions in more detail, we examined interaction heatmaps of individual chromosomes (Figure 3D). From these cis-interaction maps, we derived a 4C-style interaction plot (the interaction profile of a given locus vs. the entire chromosome) anchored at the p-arm (left) telomere of chromosome 5. In PFs and iPSCs, we did not detect prominent interactions between the p-arm (left) and q-arm (right) telomeres. Remarkably, we observed a clear increase in the normalized interaction frequency between telomeres in MNs (Figure 3D, top panel). MN-specific intra-chromosomal telomere-telomere interactions were observed for a variety of chromosomes; however, some chromosomes did not show such cis-telomeric end interactions (see below for additional analyses of these interactions during MN maturation).

### Transcriptomic and chromatin changes during early motor neuron maturation stages

We next examined how gene expression and chromatin accessibility change during the 6 weeks of motor neurons maturation in more detail. We re-differentiated two control iPSC clones (one from control-1 and one from control-2) into MNs exactly as described above. We examined genome-wide transcription and chromatin accessibility using RNA-seq and omniATAC-seq respectively at weeks 0 (W0; when post-mitotic neurons have just formed) and then at week 3 (W3), and week 6 (W6) of post-mitotic maturation (Figure 4A). For these analyses, at day 0 we isolated neuronal and non-neuronal cells from a mixed population by sorting EPCAM+ cells, which are non-neuronal, mixed epithelial-like cell lineages (EP) and NCAM+ cells, which are post-mitotic MNs.

**Figure 4.**
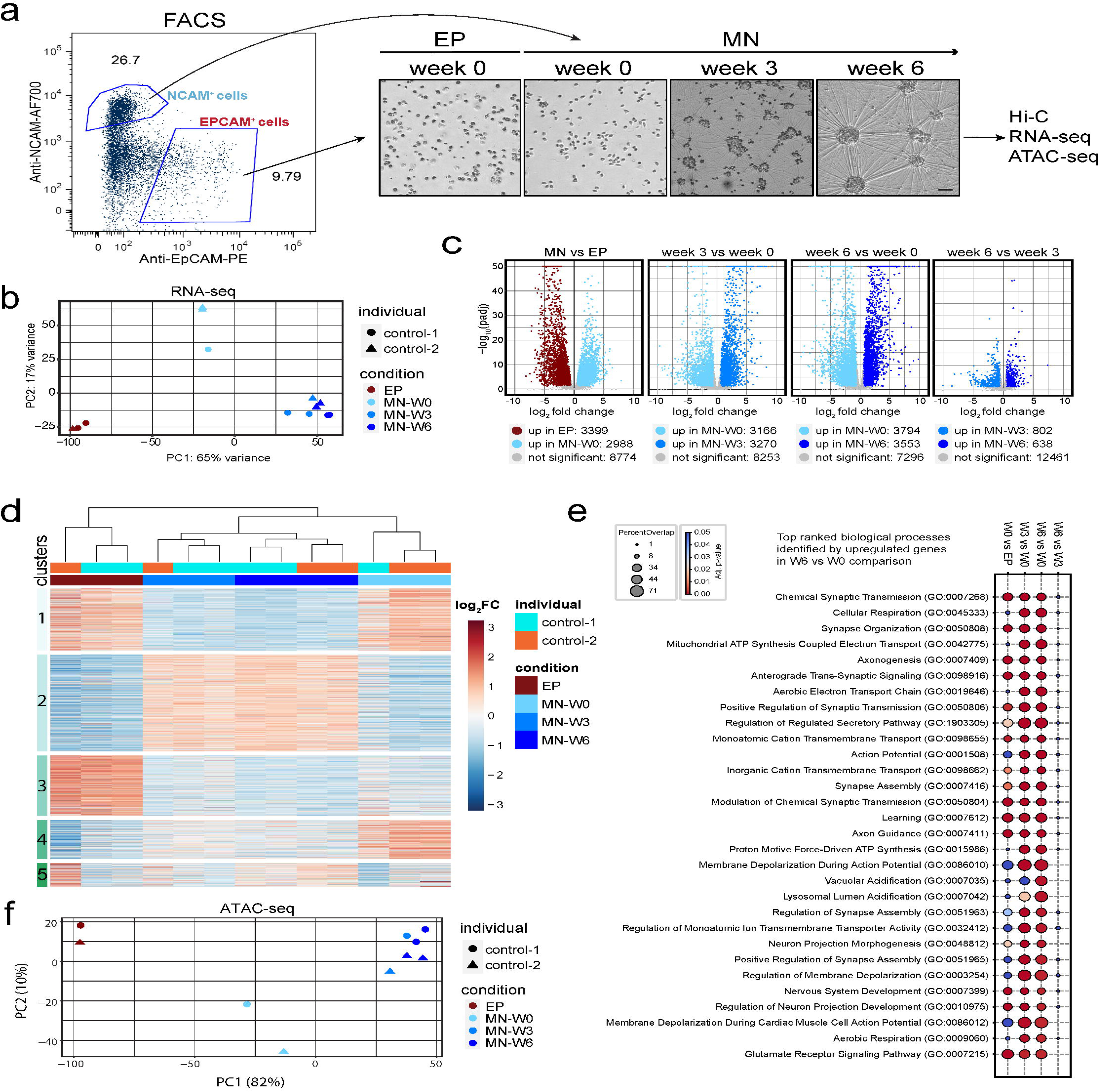
Transcriptomic and chromatin changes during early motor neuron maturation stages. a. Example of sorting settings for immature motor neurons (week 0) and EPCAM^+^ cells (EP-week 0) after 2 weeks of neural differentiation (Left panel). Brightfield images are shown during the time-course maturation of motor neurons (NCAM^+^ cells are post-mitotic immature neurons) Scale bar=50uM. b. Principal component analysis of RNA-seq samples of two control lines (Number of replicates per each time point: EP-W0=3, MN-W0=3, MN-W3=3 and MN-W6=4). c. Volcano plot distribution for RNA-seq DEseq2 comparison analyses of MN vs EP; Week 3 vs Week 0; Week 6 vs Week 0; Week 6 vs Week 3 (padj<0.05, fold change>1.5; up: upregulated). d. Heatmap shows K-mean clustering and gene expression changes of all detected genes for all time points e. Gene set enrichment analysis (GSEA) results of upregulated genes observed in the following 4 comparisons: MN-W0 vs EP-W0, MN-W3 vs MN-W0, MN-W6 vs MN-W0 and MN-W6 vs MN-W3. Percent overlap and adjusted p-values of top 30 biological processes obtained from MN-W6 vs MN-W0 comparison are plotted for every comparison (padj<0.05; Overlap percent= (Number of genes observed/ total number of genes in biological process) x100). f. Principal component analysis of the differential ATAC-seq peaks detected for two different IPSC-derived motor neuron individual lines (Number of samples per each time points: EP-W0=2, MN-W0=2, MN-W3=2 and MN-W6=4).

Principal component analysis (PCA) of all detected gene in the RNA-seq data detected three distinct groups of gene expression profiles (Figure 4B): the first contained all EP cells at week 0. The second group contained all MN-W0 cell cultures, and the third group contained all MNs at week 3 and week 6. The differences between MNs and EPs are expected, given that these are distinct cell types. The fact that the RNA-seq profiles of MNs at week 0 are distinct from those of MNs at weeks 3 and 6 of maturation shows that the transcriptome is dynamically changing during the post-mitotic maturation process, with most changes occurring during the first three weeks. In addition, the close clustering of data from different donors suggests that donor-specific differential gene expression did not play a major role.

Pair-wise comparisons of significant gene expression changes further illustrate these transcriptome dynamics (Figure 4C): MNs and EPs differ in expression for thousands of genes. Similar numbers of differentially expressed genes are detected when comparing weeks 3 or week 6 to week 0 MNs. Far fewer genes change expression in MN between week 3 and week 6.

Next, we analyzed all detected genes through k-means clustering (k=4, 5 clusters), and identified several distinct clusters associated with the different cell states (Figure 4D): cluster 1: upregulated in EP and MN at week 0; cluster 2: Maturation-specific genes (weeks 3 and 6); cluster 3: EP-specific genes; cluster 4: MN-W0-specific genes; cluster 5: a small cluster of more variable genes with difference between donors.

Finally, we performed gene ontology analysis on the upregulated genes in week 6 and compared them to MN week 0 to identify pathways activated during maturation (Xie et al., 2021). The top 30 enriched biological processes indicated that at six weeks of maturation genes related to synapse organization, neuron projection, axonogenesis, and other neural development-related processes were upregulated suggesting maturation time is critical for functional motor neurons (Figure 4E). When comparing upregulated genes between weeks 6 and 3, the top pathways became much less significant, and fewer upregulated genes were detected (Figure 4C) indicating that differentiation of the transcriptome has been mostly completed by week 3 (Figure 4C).

Next, we analyzed chromatin accessibility during maturation. PCA analysis of ATAC-seq peaks at each time point revealed a pattern consistent with the RNA-seq PCA results, with EP-W0, MN-W0, and MN-W3/MN-W6 groups showing distinct clustering (Figure 4F). In conclusion, our transcriptome and genome accessibility data show that most transcriptional and chromatin changes occur during the first three weeks of maturation. Changes in gene expression involve many hundreds of genes that encode proteins related to MN function.

### Chromatin architecture dynamics during motor neuron maturation

Above we report that mature motor neurons display increased intra-chromosomal interactions, reduced longer-range intrachromosomal (> 3Mb distance) and inter-chromosomal interactions, increased preference for homotypic compartmental interactions, and increased centromere-centromere and telomere-telomere interactions. Using the same cell preparations for immature and matured motor neurons described above, we investigated when these four changes occurred during MN maturation (Figure 5A). We find that all these characteristics are acquired after MN establishment (week 0) and during the subsequent 6-week maturation period. First, the percentage of intra-chromosomal interactions in MNs at week 0 is ∼70%, which is higher than in EPs present at that same time point (∼50%, Figure 5A and B). During MN maturation this percentage increases to ∼83% (Figure 5B). Second, the frequency of long-range intra-chromosomal interactions in MNs at week 0 is like that in EPs but is decreased by week 3, and then even more by week 6 of maturation (Figure 5C). As a result, shorter range intra-chromosomal interactions (<3 Mb) are relatively enriched in MNs compared to EPs which is also visible in Hi-C maps (Figure 5F). At that length scale loop extrusion-driven processes dominate, which can be observed in Hi-C maps as increased TADs and loops (Figure 2C), and in the shape of the derivative of *P*(*s*) (Figure 5D). Third, compartmentalization strength increases as maturation progresses, for both homotypic A-A and B-B interactions, and both in cis and in trans (Figure 5E). Fourth, at week 0, the frequency of centromere-centromere and telomere-telomere interactions in MNs are relatively weak and comparable to EPs. During maturation these interactions show remarkable increases in frequency (Figure 5A, F and G), especially for the smaller chromosomes (chromosomes 13-22). As an example, this can be readily detected using 4C-style plots anchored at the telomere of chromosome 10 (Figure 5F). We conclude that these four characteristics of chromosome and nuclear organization in mature MNs are acquired during the six-week maturation process, and not during the differentiation of iPSCs into MNs. Further, while gene expression changes mostly occur during the first 3 weeks of maturation (Figure 4C), genome folding continues to change of over the full 6 weeks.

**Figure 5.**
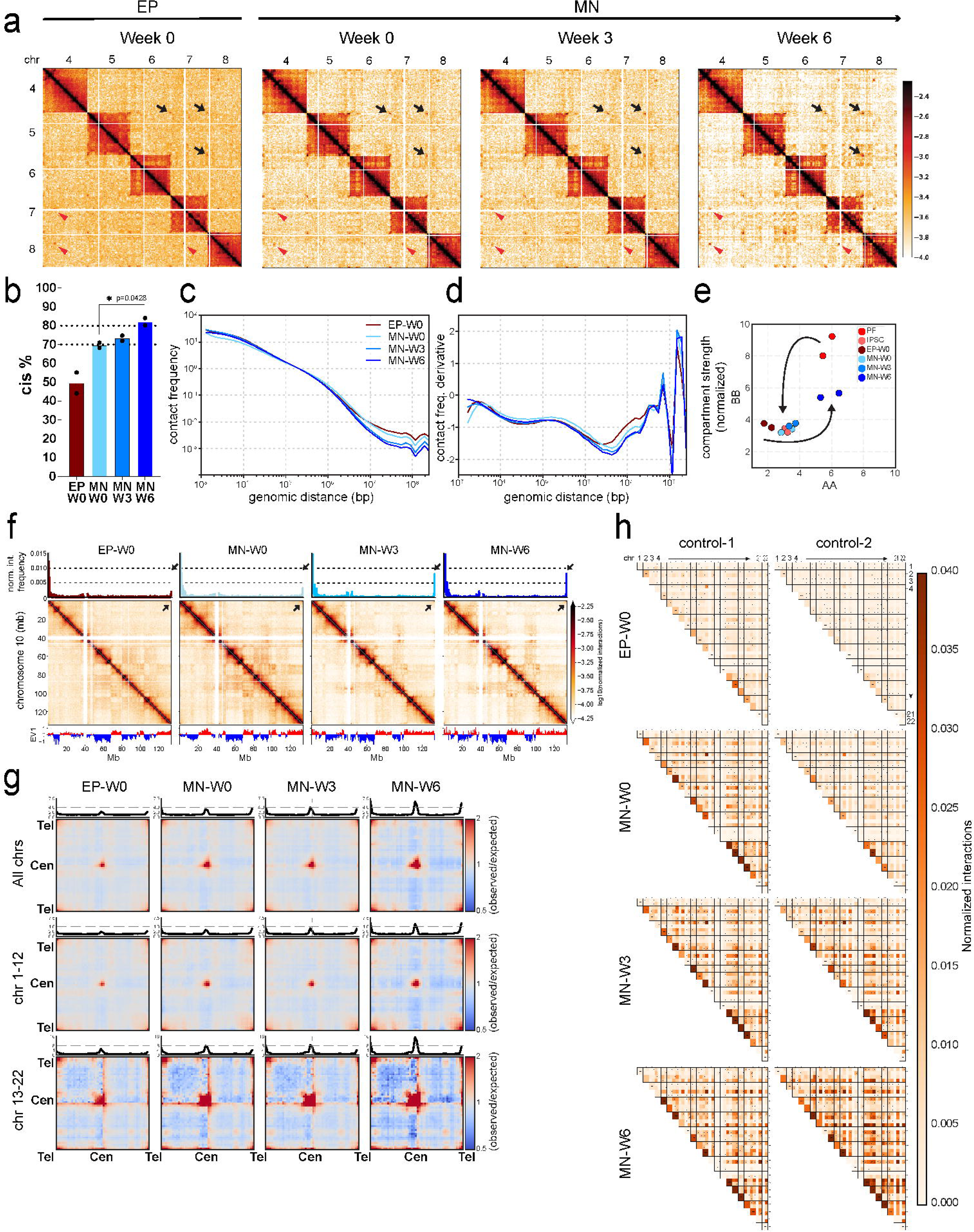
Chromatin architecture dynamics during motor neuron maturation. a. Hi-C contact matrices of trans interactions for chromosomes 4,5,6,7 and 8 in EP-W0, MN-W0, MN-W3 and MN-W6. Black arrows show telomere-telomere interactions. Red arrowheads show centromere-centromere interactions. b. Genome-wide cis interaction percentages of EP-W0, MN-W0, MN-W3 and MN-W6 (Unpaired t test with Welch’s correction was performed and only comparison of MN-W0 and MN-W6 is significant with p=0.0428). c. Genome-wide cis contact frequency plot as a function of genomic distance (1kb-∞) for EP-W0, MN-W0, MN-W3 and MN-W6. d. Derivative of contact frequency plot in panel c for EP-W0, MN-W0, MN-W3 and MN-W6. e. Compartment strengths of AA and BB obtained from the saddle plots were quantified using cis interactions for PF, IPSC, EP-W0, MN-W0, MN-W3 and MN-W6 of control #1 and #2. f. 4C plots (upper panel), Hi-C matrices (mid panel) and compartment calls (EV1: eigen value 1 obtained in PCA analysis) of cis chromatin interactions for chromosome 10 in EP-W0, MN-W0, MN-W3 and MN-W6. Black arrows show telomere-telomere interactions. g. Saddle plots of quantified genome-wide trans interactions of telomere-telomere, centromere-centromere and telomere-centromere for all chromosomes and separated chromosomes 1-12 and 13-22 for EP-W0, MN-W0, MN-W3 and MN-W6. h. Heatmap shows the normalized cis and trans interaction frequencies detected in 1 Mb region of telomeric ends among p and q arms for all autosomal chromosomes (No X, Y and M chromosomes). First column of each row indicates cis (intrachromosomal) interactions for 1 Mb region of telomeric ends (p & q arms). Squares with four sub squares demonstrate interaction frequencies of four combinations of p and q arms of two different chromosomes (top left: chr1p & chr2p; top right: chr1p & chr2q; bottom left: chr1q & chr2p; bottom right: chr1q & chr2q) for each timepoint.

We explored the increased formation of telomere-telomere interactions in more detail.

Specifically, we quantified all possible pairwise cis and trans interactions between terminal 1 Mb regions at each of the 44 telomeres of chromosomes 1-22 (Figure 5H). In EPs, telomere-telomere interactions were relatively infrequent. However, in MNs at week 0 these interactions were stronger, and they further gained in frequency during the maturation process, both in cis and in trans. Although we did observe some quantitative differences in telomere-telomere interaction frequency between two control individual MNs at week 0 (Figure 5H), by week 6 both control MNs displayed strongly increased telomeric interactions. The consistent patterning of chromosome arm interactions in MN samples suggests that these interactions are not specific to different control individuals but are uniquely and strongly detected in MNs.

The complex telomeric interaction maps for MNs at week 6 suggest that not every telomeric domain interacts with every other chromosome end. While this could be simply due to the fact that some telomeric domains may be difficult to detect with Hi-C because the presence of repetitive sequences, there are indications that there is at least some specificity in the formation of telomeric interactions. For example, for chromosome 2 we can detect both telomeres as evidenced by the fact that we observe enriched intra-chromosomal telomere-telomere interactions. Yet, the pattern of inter-chromosomal telomeric interactions for the chromosome 2 telomeres differ. Similarly, for chromosome 6 we can detect enriched intra-chromosomal telomeric interactions, but very few inter-chromosomal interactions. In contrast, the p and q arms of chromosome 17 show very strong cis interactions. The interaction frequency pattern of the p and q arm telomeres for this chromosome with other chromosome telomeres at are quite comparable. Other examples include the telomeres of chromosome 4: the p arm telomere interacts strongly with both the p and q arm telomeres of most chromosomes (Figure 5H, top left and right quadrants for every chromosome-chromosome pairwise interaction), whereas the q arm telomere does not show any enriched interactions with other chromosomes (Figure 5H, bottom left and right quadrants). In this case, we cannot rule out that the most distal telomeric region of the q arm that is detectable by Hi-C is too far from the true telomeric end to capture telomere-telomere interactions.

### *C9orf72* gene locus undergoes dynamic refolding among cell types and during maturation

Since we detected major differences in transcriptome and chromatin accessibility during maturation, we sought to examine genome folding at the scale of TADs during this process. Initially, we focused specifically on the *C9orf72* gene (chr9:27,546,546-27,573,866) locus in cells derived from healthy individuals. HRE mutations within the *C9orf72* gene cause motor neuron loss in ALS patients. Therefore, understanding how the genome folds around this key locus during maturation may provide insights into disease conditions. First, we zoomed in on the chromosome 9, 18-36 Mb region to observe how genome folding is re-arranged during reprogramming, differentiation, and maturation (Figure 6A). At the compartment level, we observed extensive changes in this region as detected by fluctuations in EV1 in accordance with changes in the checkerboard patterning in Hi-C interaction maps: in PFs the region alternates multiple times between A and B compartment status, while in iPSCs and differentiated EP and MN stages, the region is mostly in the B compartment. When we zoomed in on the *C9orf72* and *LINGO2* loci, located between 27.5-32 Mb, we observed additional changes at the domain level which reflect most likely changes at the locations of boundaries to loop extrusion (e.g., TAD boundaries), and not compartment switches given that this region is entirely embedded with a single B compartment domain (Figure 6B). In all cell types analyzed here, we observe a domain boundary at the start of the *C9orf72* locus, as reported before (Sun et al., 2018). In PFs, the rest of this region (to the right) is devoid of domain boundaries, but in iPSCs, EPs, and MNs distinct additional boundaries are clearly detectable. We also detect line-like features (green dashed boxes), that are typically ascribed to loop extrusion events anchored at a boundary (Fudenberg et al., 2017). Interestingly, while a major domain boundary is observed around position 29.8 Mb in iPSCs and MNs at week zero, this boundary disappears during MN maturation (weeks 3 and 6; black arrows). During this maturation process a new and strong boundary appears at 29.25 Mb, which coincides with the start of the *LINGO2* gene (Figure 6C; red arrows). Interestingly, there is a positive correlation between the formation of domain boundaries in weeks 3 and 6 and the expression of both genes. The expression of *C9orf72* increased more than 2-fold between week 0 and week 6. The expression of *LINGO2* was barely detectable in week 0 MN and EP lines. After 3 weeks of maturation, expression levels increased, but by week 6, levels were reduced by 2-fold (Figure 6D). This activation is also reflected in changes in the value of EV1 between the 27.5-29.5 Mb regions among cell types, with a tendency to increase EV1 so that it is less negative as motor neurons mature (Figure 6E and 6F). Further, we note that an additional domain boundary appears in MNs approximately 250 kb upstream of the *C9orf72* gene. This boundary is relatively weak at week 0 and then becomes more pronounced during maturation as seen in the insulation profile (Figure 6C). This boundary coincides with an ATAC-seq peak that appears around week three of maturation (Figure 6F-red arrow).

**Figure 6.**
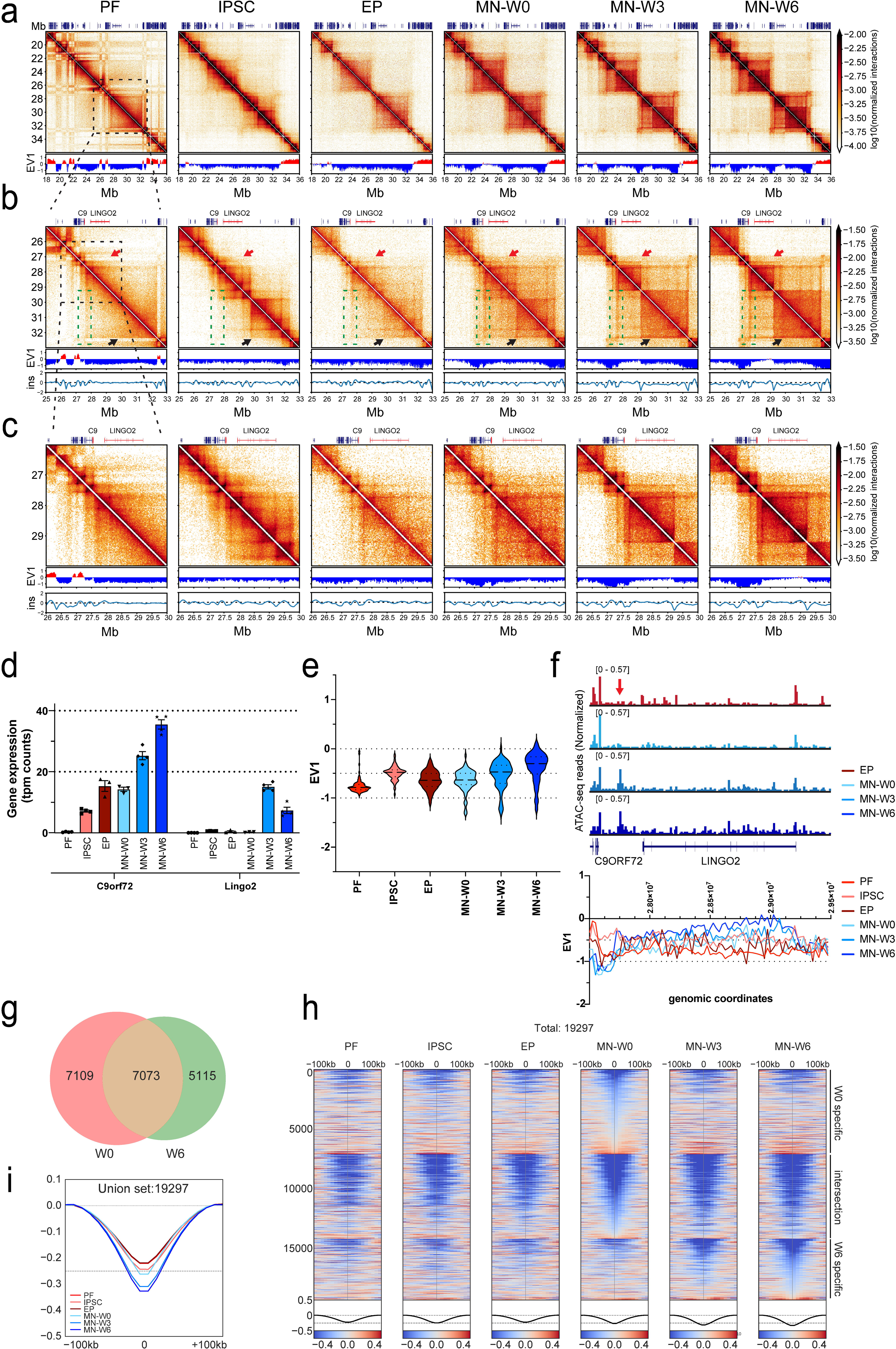
C9orf72 gene loci undergo dynamic rearrangements among cell types and during maturation. a. Hi-C contact matrices of chromosome 9: 18-36 Mb in PF, IPSC, EP-W0, MN-W0, MN-W3 and MN-W6. lower panel: compartment calls obtained from EV1 (100kb bin size). b. Hi-C contact matrices of chromosome 9: 25-33 Mb in PF, IPSC, EP-W0, MN-W0, MN-W3 and MN-W6. Mid panel: compartment calls obtained from EV1 (25kb bin size). Bottom panel: insulation scores (ins) for TAD boundary calls (25kb bin size; 250kb window size); C9: *C9orf72* gene. c. Hi-C contact matrices of chromosome 9: 26-30 Mb in PF, IPSC, EP-W0, MN-W0, MN-W3 and MN-W6. Mid panel: compartment calls obtained from EV1 (25kb bin size). Bottom panel: insulation scores (ins) for TAD boundary calls (25kb bin size; 250kb window size); C9: *C9orf72* gene. d. Gene expression profiles of the *C9ORF72* and *LINGO2* genes that are obtained from RNA-seq for each condition (tpm: transcript per million). e. Violin plot distribution of Eigen Value 1 (EV1) at 25kb resolution between 27.5-29.5 Mb region among all cell-types. f. Normalized ATAC-seq distribution of EP-W0, MN-W0, MN-W3 and MN-W6 (top panel) and EV1 distribution among all cell-types between 27.5-29.5 Mb locus in the *C9ORF72* and *LINGO2* gene regions (bottom panel; both genes are transcribed in the minus direction-right to left). g. Venn diagram of strong boundaries compared between week 0 and week 6 (W0= week 0 specific: 7109; W0&W6= common: 7073; W6= week 6 specific: 5115). h. Stack-up plots of insulation scores detected at the strong boundaries with flanking 100kb from each side. W0-specific, intersection, and W6-specific strong boundaries detected in figure 6g are used. i. Aggregated insulation score pileup distributions of strong boundaries detected in the union set of weeks 0 and 6 among various cell types (PF: primary fibroblast; IPSC: induced pluripotent stem cell; EP: EPCAM^+^ cells; MN: motor neurons).

In summary, the genomic region containing *C9orf72* and *LINGO2* displays dynamic changes in chromatin organization during cell differentiation, as well as during post-mitotic MN maturation. New domain boundaries form at promoters and other open chromatin sites as detected by ATAC-seq.

### Strengthening of TAD boundaries during motor neuron maturation

Next, we analyzed TAD boundary positions and strengths genome-wide during maturation. We identified boundaries in EP-W0, MN-W0, MN-W3, and MN-W6 using a threshold for strong boundaries (boundary strength > 0.1; see Methods). MN-W0 and EP-W0 cells showed the highest number of boundary calls. By week 6, the total number of boundaries was reduced by ∼20% (Figure S6A).

We determined the union set of all boundaries in immature and mature MNs (Week 0 vs Week 6 union set: 19,297 boundaries). Among these, 7,109 were specific to week 0, and 5,115 were specific to week 6. The remaining 7,073 boundaries were observed in both datasets (Figure 6G). We aggregated all insulation scores at the boundaries and their flanking 200 kb regions, categorizing them into three groups: week 0-specific (W0), common (intersection of W0 & W6), and week 6-specific (W6) (Figure 6H). As expected, most of the week 0-specific boundaries were lost by the week 3 and 6 time points. However, most week 6-specific boundaries were also present at week 3, indicating that a significant number of TAD boundaries already formed by week 3 and became stronger over the 6-week maturation period. The intersecting set of common boundaries seen in week 0 and week 6 were also detected in non-neuronal cells, suggesting that

these boundaries are maturation-independent and conserved across different cell types. We then aggregated all insulation scores around the boundaries obtained from the union set for PF to MN-W6. Interestingly, MN-W3 and MN-W6 showed the strongest insulation scores, indicating that overall boundaries were stronger in mature MNs (week 6) than in immature MNs (week 0) and other cell types (Figure 6I). We conclude that boundary formation is dynamically changing during maturation, while boundary strength increases.

### Impact of *C9orf72* HRE on Chromatin Folding in ALS Patient-Derived Cells

Our studies in control cells above show that during differentiation and MN maturation chromosome folding, nuclear organization, chromatin accessibility and gene expression show large scale changes. We next set out to investigate whether these phenomena are affected in cells derived from ALS patients with *C9orf72* HRE mutations (Figure S7A-C).

We first focused on chromosome conformational dynamics (Figure 7A, 7E and S7D). We performed Hi-C on fibroblasts, iPSCs, and iPSC-derived motor neurons (six-week matured) from ALS patients with *C9orf72* HRE mutations. As described above, in cells from normal donors, we observed key changes in chromosome folding: 1) changes in compartmentalization patterns and strength during reprogramming and differentiation; 2) Reduction in long-range intra-chromosomal and inter-chromosomal interactions; 3) Increased centromere-centromere and telomere-telomere interactions during maturation; and 4) changes in domain boundary positioning and strength during maturation. We quantified these features in cells derived from ALS patients carrying *C9orf72* HRE mutations. First, visual inspection of intra-chromosomal Hi-C maps shows no obvious differences for PFs and iPSCs for cells derived from normal individuals or ALS patients (Figure S7D, first three columns). Interestingly, while in mature MNs from normal donors we observed an increase in compartmentalization strength (Figure 2E), in MNs from ALS patients this did not occur or to a reduced extend (Figure S7F and G). This is visible in Hi-C maps by strong long-range compartmental intra-chromosomal interactions observed in control motor neurons (interaction of the 10 Mb and 56 Mb regions in MN1 and MN2, Figure S7D, arrows)) and partial or near complete loss in these interactions in Hi-C maps for MN clones from ALS cases (Figure S7E (arrows), columns 4 and 5). Genome-wide saddle plot analysis confirmed this change in compartmentalization strength as compared to matured MNs from controls (Figure S7F and G). Although all four MN clones showed reduced compartmentalization strength, there is clearly variation between them. We note that the ALS patient-derived MN clone that showed the smallest reduction in compartmentalization strength (C9-ALS-2 MN clone 2), was derived from an iPSC clone that already showed relatively strong compartmentalization strength and high intra-chromosomal interaction frequency (Figure 7C, red data points) compared to other iPSC clones, suggesting that variation in reprogramming could contribute to the observed variation in MN Hi-C data (see discussion).

**Figure 7.**
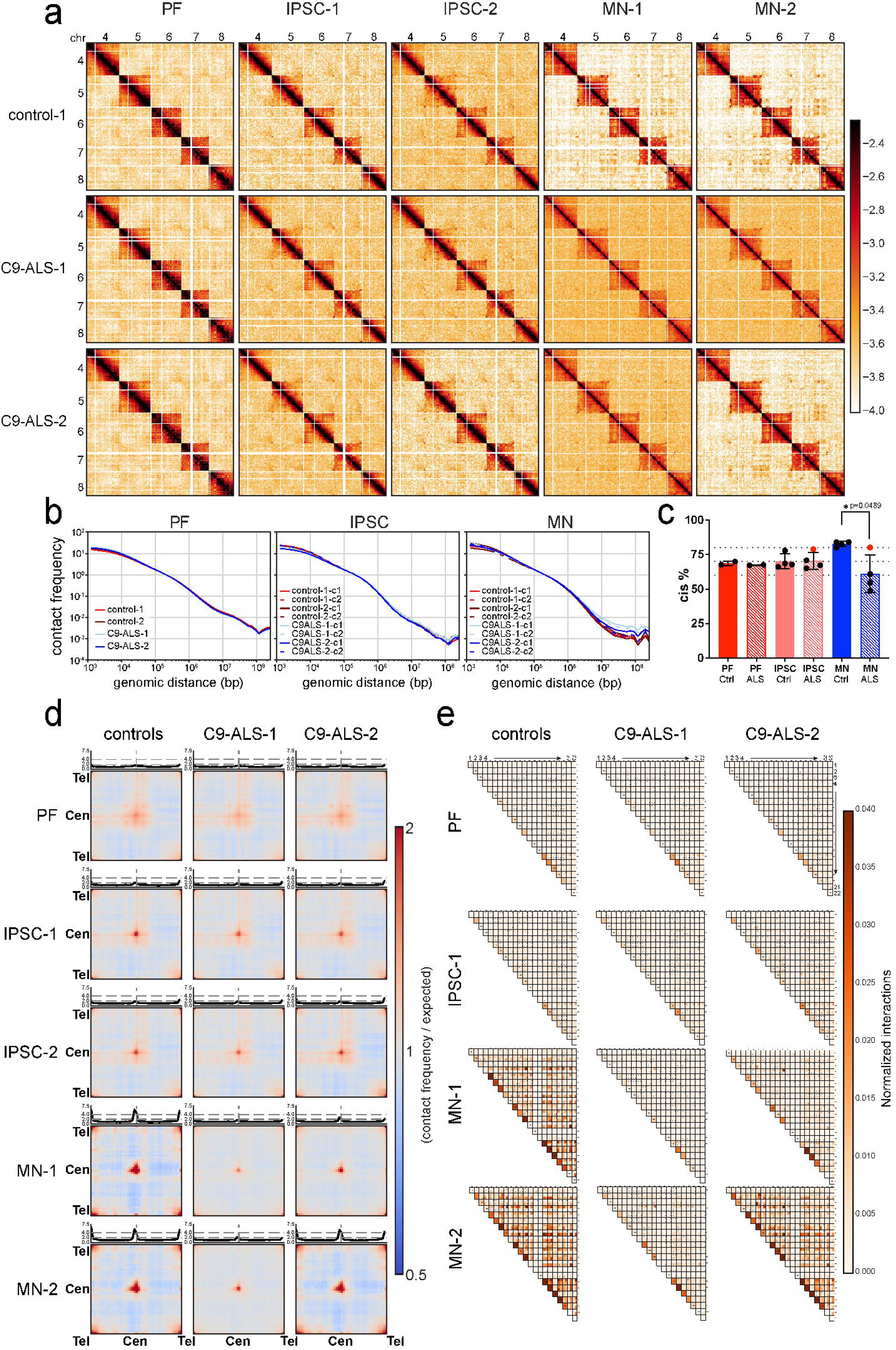
Impact of C9orf72 HRE on Chromatin Folding in ALS Patient-Derived Cells. a. Hi-C contact matrices of trans interactions of chromosomes 4,5,6,7 and 8. PF, two IPSCs and two MNs of control 1 and two ALS patients. b. Genome-wide contact frequency *P* as a function of genomic distance *s* between interaction pairs is plotted for PFs, IPSCs and MNs of controls and ALS patients. c. Genome-wide intra-chromosomal (cis) interaction percentages in PFs, IPSCs and MNs of controls and ALS patients (Unpaired t test with Welch’s correction was performed and only comparison of MNs from controls and ALS patients is significant with p=0.0489). d. Saddle plots of quantified genome-wide trans interactions of telomere-telomere, centromere-centromere and telomere-centromere of all chromosomes for PFs, IPSCs and MNs of controls and ALS patients. e. Heatmap shows the normalized cis and trans interaction frequencies detected in 1 Mb region of telomeric ends among p and q arms for all autosomal chromosomes (No X, Y and M chromosomes). First column of each row indicates cis (intrachromosomal) interactions for 1 Mb region of telomeric ends (p & q arms). Squares with four sub squares demonstrate interaction frequencies of four combinations of p and q arms of two different chromosomes (top left: chr1p & chr2p; top right: chr1p & chr2q; bottom left: chr1q & chr2p; bottom right: chr1q & chr2q) for PFs, IPSCs and MNs of controls and ALS patients.

Second, we analyzed long-range intra-chromosomal and inter-chromosomal interaction frequencies. For instance, Figure 7A shows inter-chromosomal interactions for chromosomes 4 to 8. Among all individuals (both cases and controls), PF and iPSC clones demonstrated very similar interaction frequency patterns within their respective cell types (Figure 7A, columns 1, 2, and 3). However, in MN clones, we found that overall levels of inter-chromosomal interactions were increased in three MN clones of C9-ALS cases as compared to controls (Figure 7A, C).

One MN clone of C9-ALS-2 (c2) showed a level of inter-chromosomal interactions similar to control clones (Figure 7A, columns 4 and 5). Inter-chromosomal compartmentalization strength was reduced in all ALS-derived MNs as compared to controls (Figure S7F and G), including for the one MN clone that showed no change in the level of inter-chromosomal interactions. These results show that in ALS-derived matured MNs, the reduction in inter-chromosomal interaction frequencies and increased compartmentalization strength that normally occurs during MN maturation does not occur. Again, we note that there is variation among the four ALS-derived MN clones, as there is among the controls.

We inspected *P*(*s*) plots to assess long-range intra-chromosomal interactions (Figure 7B). *P*(*s*) for Hi-C data obtained from PFs and iPSCs derived from normal and ALS donors were indistinguishable (Figure 7B). However, differences were apparent when *P*(*s*) plots for MNs were compared. For MNs derived from normal donors we observed a reduction in long-range interactions (>2 Mb). This reduction was not observed in both MN clones derived from ALS patient 1 (C9ALS-1 c1 and C9ALS-1 c2), and in one of two clones derived from ALS patient 2 (C9ALS-2 c1 and C9ALS-2 c2). Similarly, the increased level of intra-chromosomal interactions detected in control MNs as compared to PFs and iPSCs, was not observed for three of four MN clones derived from ALS patients (Figure 7C). The one ALS-derived MN clone that showed the least change in intra-chromosomal interaction frequencies again corresponded to the same clone that showed the smallest (but detectable) defect in compartmentalization strength (above).

Third, we investigated the frequencies of centromere-centromere and telomere-telomere interactions. These increased during maturation of MNs derived from normal donors (Figure 3C). In PFs and iPSCs derived from ALS patients, we did not detect differences in these interactions compared to PFs and iPSCs derived from controls (Figure 7D and E). However, for three out of four MN clones derived from ALS patients we did not observe an increase in centromere-centromere and telomere-telomere interactions as was seen in controls. The fourth clone showed, as above, a considerably smaller but still detectable defect in increase in centromere-centromere and telomere-telomere interactions.

We determined how p and q arm telomeric interactions are altered in C9-ALS patients compared to controls. We plotted pairwise inter-chromosomal telomeric end interactions for all four possible combinations among PF, iPSC, and MNs, as we did above (Figure 3C; Figure 7E). As expected, all PF lines including controls and cases, demonstrated similarly weak inter-chromosomal interactions at telomeric ends (normalized contact frequency ≤0.001). In MNs from ALS patients, for three out of four clones, we did not observe the increased telomere-telomere interactions that is observed in controls. Intra-chromosomal telomeric interactions were also relatively weak overall, though slightly increased for smaller chromosomes. We again note some variation in telomere-telomere interactions between clones with the second MN clone from patient C9-ALS-2 again being less affected. In summary, in MNs from ALS patients, major changes in large-scale chromosome folding normally observed during maturation were not detected or reduced in effect size.

### *C9orf72* HRE does not lead to major changes in chromosome conformation around the *C9orf72* locus

We next examined Hi-C data for the *C9orf72* locus and surrounding chromatin to understand whether the HRE mutation impacted chromatin architecture around the gene. First, we analyzed Hi-C maps and insulation scores along the chromosome 9, 26-30 Mb region (Figure S7H). As described above in control cells, domain boundaries throughout this region show changes as cells differentiate and mature into MNs (Figure 6A-C). These changes occurred normally in cells from ALS patients (Figure S7H). The *C9orf72* gene is located near a TAD boundary at ∼27.5 Mb. In PF lines, no alterations were observed between controls and cases in terms of Hi-C maps (Figure S7F). Similarly, in MNs boundaries surrounding the *C9orf72* and *LINGO2* gene showed a nearly identical pattern across controls and ALS patients. We conclude that cell type-specific changes in Hi-C and insulation patterns occur normally in cells obtained from ALS patient cells.

Finally, we analyzed insulation at boundaries genome wide. Using the union set of boundaries identified for control MNs (Figure 6G), we quantified insulation at these locations in cases and controls (Figure S7I, J and K). While we detected some variation between clones, we did not observe consistent changes specific to MNs derived from ALS patients. We conclude that boundary formation and strength is not majorly affected in ALS.

### Transcriptome and genome accessibility impacts of *C9orf72* HRE Mutation in ALS Motor Neurons

We performed total RNA-seq to determine whether the HRE mutation has a transcriptome-wide impact in iPSCs and MNs. We analyzed four datasets for ALS-derived cells and four datasets from controls (two donors, two clones each), and determined changes in gene expression using DEseq2 (Love et al., 2014). The gene expression profile in iPSCs derived from C9-ALS patients did not show major differences compared to controls (40 genes are downregulated and 30 genes upregulated in ALS-derived cells; Figure 8A). When we compared the transcriptomes of C9-ALS-derived mature MNs (W6) to control MNs (W6), we observed more differentially expressed genes: we detected 372 downregulated and 299 upregulated genes in C9-ALS motor neurons (Figure 8B). Further, when we compared the transcriptomes of C9-ALS-derived mature MNs (W6) to control MNs at week 0, we find that large numbers of genes change expression, in concordance with the comparison of transcriptomes of control MNs (W6) to control MNs (W0) (Figure 4C). Combined these results show that as C9-ALS cells differentiate from iPSCs to matured MNs, thousands of genes change their transcriptional status and the final mature MNs resemble those of controls. This is confirmed by GO-term analysis: gene categories upregulated in control mature MNs during maturation include categories related to neuronal functions, synapse formation, and axonogenesis which are also significantly upregulated in C9-ALS MNs (Figure 8C, compare column 4 and 9). This result shows that C9-ALS MNs mature to a considerable extent in terms of their gene expression profile.

**Figure 8.**
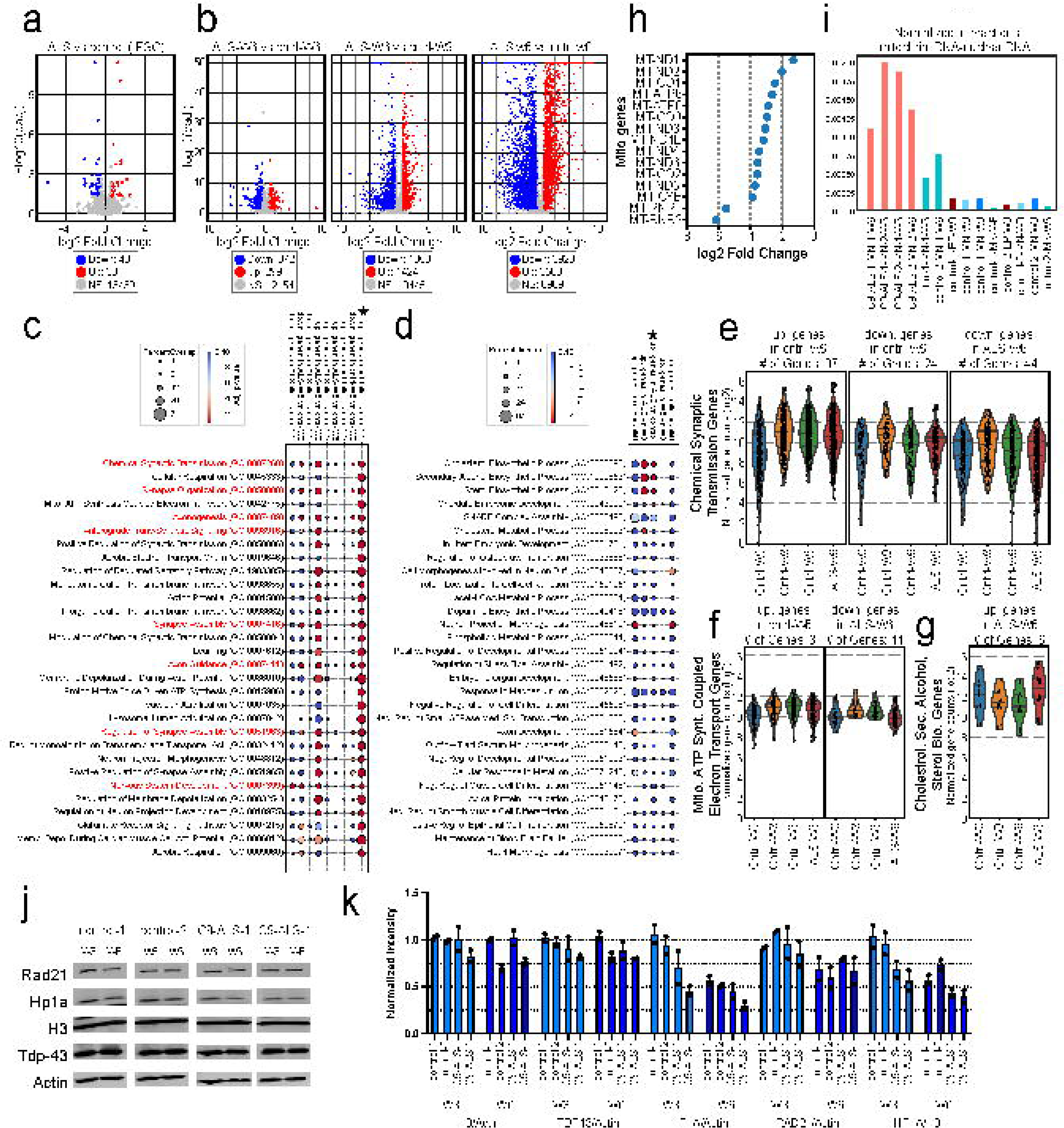
Transcriptome impacts of C9orf72 HRE Mutation in ALS Motor Neurons. a. Volcano plot of DEseq2 comparison for ALS vs Control IPSC lines (padj<0.05, fold change>1.5; up: upregulated, down: downregulated, NS: not significant). b. Volcano plots of DEseq2 comparisons for ALS-W6 vs Control-W6, ALS-W6 vs Control-W3, ALS-W6 vs Control-W0 MNs (padj<0.05, fold change>1.5; up: upregulated, down: downregulated, NS: not significant). c. Biological processes (gene ontology) are ranked according to the top 30 lowest p-values detected in up. genes in cntrl-W6 vs cntrl-W0 (starred). Percent overlap and p-values for each biological process observed in different pairwise comparisons are plotted for the following 9 columns (left to right): down. genes in ALS-W6 vs cntrl-W0, down. genes in ALS-W6 vs cntrl-W3, down. genes in ALS-W6 vs cntrl-W6, up. genes in ALS-W6 vs cntrl-W0, up. genes in ALS-W6 vs cntrl-W3, up. genes in ALS-W6 vs cntrl-W6, up. genes in cntrl-W6 vs cntrl-W3, down. genes in cntrl-W6 vs cntrl-W3, up. genes in cntrl-W6 vs cntrl-W0 (Percent overlap= (Number of genes observed/ total number of genes in biological process) x100). d. Biological processes (gene ontology) are ranked according to the top 30 lowest p-values detected in up. genes in ALS-W6 vs cntrl-W6 (starred). Percent overlap and p-values for each biological process observed in different pairwise comparisons are plotted for the following 5 columns (left to right): up. genes in ALS-W6 vs cntrl-W0, up. genes in ALS-W6 vs cntrl-W3, up. genes in ALS-W6 vs cntrl-W6, up. genes in cntrl-W6 vs cntrl-W3, up. genes in cntrl-W6 vs cntrl-W0 (Percent overlap= (Number of genes observed/ total number of genes in biological process) x100). e. Expression levels of genes associated with the chemical synaptic transmission biological process were plotted across four conditions: cntrl-W0, cntrl-W3, cntrl-W6, and ALS-W6. The panels represent three comparisons. Left panel: up. genes in cntrl-W6 vs cntrl-W0 (# of Genes: 87); mid panel: down. genes in cntrl-W6 vs cntrl-W3 (# of Genes: 24); right panel: down. genes in ALS-W6 vs cntrl-W3 (# of Genes: 44). f. Expression levels of genes associated with the mitochondrial ATP synthesis coupled electron transport biological process were plotted across four conditions: cntrl-W0, cntrl-W3, cntrl-W6, and ALS-W6. The panels represent two comparisons. Left panel: up. genes in cntrl-W6 vs cntrl-W0 (# of Genes: 31); right panel: down. genes in ALS-W6 vs cntrl-W3 (# of Genes: 11) g. Expression levels of genes associated with the cholesterol, secondary alcohol and sterol biosynthetic processes found in up. genes in ALS-W6 vs cntrl-W6 comparison were plotted across four conditions: cntrl-W0, cntrl-W3, cntrl-W6, and ALS-W6. h. Fold change expression levels of mitochondrial genome-encoded genes identified among the downregulated genes in the ALS-W6 vs cntrl-W6 comparison are shown. i. Normalized pairwise interactions of mitochondrial genomic DNA and nuclear genomic DNA as detected by Hi-C data, were plotted for time-course and MN-W6 samples. j. Representative western blot images of week 3 and week 6 motor neurons (MNs) from control-1, control-2, and two distinct clones of C9-ALS-1 were analyzed using antibodies against Rad21, HP1a, H3, TDP-43, and Actin. k. Protein levels of Rad21, HP1a, TDP-43, and H3 were quantified with intensities normalized to Actin or H3. Fold changes were then plotted.

However, there are also some notable differences. Several of the gene categories upregulated in control MNs, are not significantly upregulated in C9-ALS MNs (W6), e.g. many related to cellular respiration, mitochondrial ATP synthesis coupled electron transport, and proton motive force-driven ATP-synthesis. Furthermore, we investigated gene expression levels in control-W0, control-W3, control-W6 and C9-ALS-W6 MNs that are involved in significantly upregulated or downregulated biological processes (Figure 8C) such as chemical synaptic transmission (Figure 8E), synapse organization, axonogenesis and anterograde trans-synaptic signaling processes (Figure S8A-C). For example, we compared upregulated or downregulated genes that are identified in pairwise comparisons done in figure 8C: columns 9, 8 and 2 for chemical synaptic transmission genes, respectively (Figure 8E). Expression levels of the genes detected in control-W6 vs control-W0 comparison (column 9) are also at similar levels for control-W3 and C9-ALS-W6. Likewise, expression levels of genes detected in columns 8 and 2, show similar expression profiles in control-W6 and ALS-W6 conditions. This expression pattern is also observed in other highlighted processes (Figure S8A-C) indicating that C9-ALS-W6 MNs have similar expression profiles for the genes found in these biological processes. However, when we compared gene expressions of non-significant biological processes such as mitochondrial ATP synthesis coupled electron transport, seen in upregulated genes in ALS-W6 vs control-W0 comparison (column 4), we observed reduced gene expression profiles in ALS-W6 compared to control-W3 and control-W6 (Figure 8F).

Importantly, as shown in Figure 8B, C9-ALS MNs (W6) do show differentially expressed genes as compared to control MNs (W6). We did not detect any significant enrichment for specific GO terms in the small set of genes that are downregulated in C9-ALS MNs compared to control MNs. However, the upregulated gene set is significantly enriched in genes involved in cholesterol, secondary alcohol, and sterol synthesis (Figure 8D and G). In control MNs, these genes are downregulated throughout the 6-week maturation period. In C9-ALS MNs this downregulation occurs to a lesser extent.

Taken together, transcriptome analysis shows that C9-ALS MNs neurons mature as reflected in the global changes in their gene expression profile. Compared to control MNs, we observe reduced expression of genes involved in cellular respiration and increased expression in genes involved in cholesterol, alcohol, and sterol synthesis.

Since we observed defects in expression of genes involved mitochondrial activity, respiration and ATP synthesis processes, we specifically investigated expression of mitochondrially encoded genes (Figure 8H). We find that there is a dramatic reduction for fifteen mtDNA protein coding genes suggesting that there is a mitochondrial dysfunction or reduced number of mitochondria in ALS-W6. Independently, we found in our Hi-C datasets elevated interactions between the mitochondrial and nuclear genome in C9-ALS MNs as compared to controls (Figure 8I). These interactions are background signals for Hi-C, but their elevated levels in C9-ALS MNs indicate potential loss of mitochondrial integrity. Combined these lines of evidence indicate defects in structure and function of mitochondria in C9-ALS MNs.

In addition, we analyzed genome-wide transcription factor (TF) binding dynamics using foot printing analysis of ATAC-seq data from control MNs (W0, W3, and W6), and C9-ALS MNs (W6). We used TOBIAS (Bentsen et al., 2020) to determine “binding score” for each transcription factor (TF) in each sample. We then calculated pairwise comparisons of TF binding activity for all TF motifs during the maturation process and the disease state of MNs. Figure S8G shows a heatmap of binding score differences for TFs showing the top up-and down-regulated differential binding activity during control MN maturation. Pairwise comparisons revealed sets of TF binding motifs that become more or less accessible during the normal maturation process (MN-W0 vs MN-W3, MN-W0 vs MN-W6, and MN-W3 vs MN-W6). The same patterns of increased and decreased accessibility for these TFs were observed when we compared ATAC-seq data from C9-ALS MNs (W6) to control MNs (W6). This shows that ATACseq profiles obtained with ALS-derived motor neurons at week 6 differ from those obtained with control mature motor neurons. Further, they differ in the same way from control motor neurons as control week 3 differs from data obtained from control motor neurons at week 6. This is also illustrated by the fact that TF binding site accessibilities of week 6 ALS-derived MNs resemble that of control MNs at week 3. We conclude that TF activity and chromatin accessibility in week 6 motor neurons fom C9-ALS MNs resembles the less mature state of control MNs at week 3 of maturation.

Finally, our MN Hi-C results showed that chromatin architecture is mainly affected in heterochromatic regions, e.g., regions near centromeres and telomers. However, we cannot conclude which pathological feature of the *C9orf7*2 gene (loss of function of the *C9orf72* gene, RNA foci, or DPR aggregate toxicity) is the leading cause of this phenotype. A study on DPR toxicity showed that overexpression of one of the most toxic DPRs, proline-arginine (PR), in mice disrupts HP1α liquid phases due to decreased expression of the HP1α gene and dissociation from chromatin, causing de-repression of heterochromatin and leading to genome instability (Zhang et al., 2019b). We re-differentiated and matured additional MNs from two control and two clones of the one C9-ALS line and determined the expression levels of key nuclear proteins including Rad21, HP1α, and TDP-43 using western blotting. Interestingly, we found that Rad21, and TDP-43 levels were slightly reduced when comparing week 3 to week 6 timepoints.

However, HP1α levels were reduced by half at week 6 compared to week 3. Possibly, the reduction of HP1α protein levels and/or function contributes to the reduced levels of centromeric and telomeric interations we observed in the Hi-C maps in C-ALS-derived MNs (Figure 8J and K).

## Discussion

We show that the genome undergoes large-scale changes in folding, accessibility, and gene transcription as cells reprogram, differentiate into motor neurons and then mature during up to six-week post-mitotic neuronal maturation. As expected, as cells reprogram and differentiate, thousands of genes change expression, and genome accessibility and conformation (e.g., compartmentalization) changes. Here, we focus mostly on the process of post-mitotic maturation. During this process thousands of genes change expression, especially during the first three weeks, making matured (week 6) motor neurons and early motor neurons (week 0) as distinct in their transcriptome as different cell types. The organization of chromatin and chromosomes, (e.g., the clustering of centromeres and telomeres) continues to change throughout the six weeks, even after the transcriptome is largely stabilized. This suggests that a slow biophysical process drives centromere and telomere clustering. This process is defective in neurons derived from ALS patients carrying *C9orf72* HRE mutations.

During maturation, thousands of genes become activated and repressed. Genes that are upregulated are involved in motor neuron function including synapse organization, transmission, and axon guidance as well as energy metabolism such as cellular respiration, electron chain transport and ATP synthesis. After three weeks of maturation the transcriptome stabilizes. Morphologically, cells are clustered with long axons protruding, and they are fully active as neurons.

Chromosome conformation, chromatin accessibility and nuclear organization continue to evolve over the six-week maturation period studied here. We observed a change in the pattern of long-range chromatin interactions: there is a reduction in inter-chromosomal interactions, very long-range intra-chromosomal interactions (loci separated by >1-2Mb), and shorter-range interactions start to dominate. Similar results were found in other studies (Heffel et al., 2024; Hu et al., 2021; Pletenev et al., 2024). These short-range interactions are in part driven by cohesin-mediated loop extrusion, suggesting that loop extrusion is a dominant driver of chromosome folding in neurons. Consistent with this, we observe increased in TAD boundary strength, and the shape of the derivative of *P*(*s*) changes with a steeper slope at genomic distances of several Mb. This latter feature has been linked to increased cohesin loop density (Gassler et al., 2017).

Increased loop extrusion often counteracts the process of chromosome compartmentalization (Haarhuis et al., 2017), but this is not what we observe in mature motor neurons: while the absolute frequency on long-range (>1-2 Mb) interactions and inter-chromosomal interactions decrease during maturation, compartmentalization strength, i.e., the preference for A loci to interact with other A loci and B loci to interact with other B loci, increases. We observe highly localized long-range interactions, both in cis and in trans, often between active genes. Such clustering, especially between long genes, has been observed in neurons before (Takei et al., 2025). These clusters appear distinct from nuclear speckles, and what drives their formation is not known. In other studies, it was found that such focal interactions can also involve polycomb-bound loci (Pletenev et al., 2024). It is likely that this is also the case in the matured neurons we study here, though we do not currently have polycomb-related chromatin data for these cells.

In a recent study of chromosome organization in the brain, it was also reported that in neurons, as compared to non-neuronal cells, shorter range interactions dominate over longer range interactions (Heffel et al., 2024). However, in that study weaker compartmentalization was observed. It is not clear how this difference with our results can be explained. We do note that identifying compartment profiles along chromosomes in neurons using the widely used eigenvector decomposition method (Imakaev et al., 2012) is not trivial because long-range interactions are much reduced in overall frequency, and as a result the largest eigenvector may not always correspond to the compartment profile. We manually curated all compartment profiles to avoid misassignments of compartment status, which in turn would result in underestimating compartmentalization strength.

We observed continuous increases in interactions between centromeres, and between telomeres as cells mature throughout the maturation period. This suggests a reorganization of relative positions of chromosome territories. Such arrangement resembles a “Rabl” orientation, but while the Rabl arrangement is typically associated with a recent anaphase, this cannot be the case here given that these cells are non-dividing. It is likely that these interactions are driven by heterochromatic associations. Previously, it was found that when heterochromatin is artificially detached from the nuclear envelope, the nucleus slowly over several weeks inverts so that all heterochromatin coalesce at the center of the nucleus (Solovei et al., 2013). This process has been proposed to be driven by biophysical attractions between heterochromatic loci, e.g., through HP1 proteins and tends to be slow as it involved large-scale movement of chromosomes (Falk et al., 2019). It is intriguing that centromeres cluster with centromeres, but not with telomeres. The composition of heterochromatin at these loci may be distinct, or these clusters may occur at different sub-nuclear locations. The functional relevance of this organization is not known, but it is interesting that this process is defective in neurons derived from ALS patients carrying *C9orf72* HREs. A recent study found that expression of poly(PR), a proline-arginine (PR) dipeptide repeat protein expressed from the expanded repeat in the *C9orf72* gene, localizes to heterochromatin and leads to reduced HP1a expression and activity (Zhang et al., 2019b).

Fibroblasts and iPSCs derived from ALS patients carrying a hexanucleotide repeat expansion in the *C9orf72* locus do not appear distinct from cells derived from controls: they show prominent compartmentalization in fibroblasts and weaker compartmentalization in iPSCs as reported previously (Akgol Oksuz et al., 2021), and the difference in transcriptomes between patients and controls is minimal. However, we observed differences in matured motor neurons. Motor neurons from ALS patients do not show the same changes in chromosome conformation and nuclear organization as controls. Centromere clustering and telomere clustering is weaker, compartmentalization is weaker, and long-range intra-chromosomal interactions and inter-chromosomal interactions do not go down in frequency as occurs during maturation of control motor neurons. It is intriguing that HP1 proteins, that may be involved in mediating such heterochromatic interactions are expressed at lower levels in ALS neurons. Other studies also reported defects in heterochromatin maintenance in ALS (Jury et al., 2020; Zhang et al., 2019b). Finally, chromatin accessibility of week 6 neurons from ALS patients resembles control motor neurons at week 3 based on TF footprinting. These observations suggest that maturation of the chromosome state is not occurring normally.

During maturation, motor neurons from ALS patients display similar large-scale changes in the transcriptome as control motor neurons. However, we noted some gene expression changes of interest. For instance, genes involved in sterol and alcohol biosynthesis are not down regulated to the same extent as in controls. Inappropriate expression of genes involved in these processes was recently described for an ALS mouse model with the *SOD1* mutation (Dominov et al., 2023). Second, we noticed that genes involved in cellular respiration, electron transport chain, and mitochondrially encoded genes are downregulated. This suggests a mitochondrial defect which is in concordance with previous findings that early mitochondrial dysfunction, endoplasmic reticulum stress combined with impaired mitochondrial Ca² buffering and suppression of local translation of nuclear-encoded mitochondrial proteins are observed in IPS-derived motor neurons with various ALS causing mutations (Altman et al., 2021; Dafinca et al., 2020; Pilotto et al., 2025; Schweingruber et al., 2025). Interestingly, we observed increased nuclear genome-mitochondrial genome interactions in Hi-C data obtained from ALS neurons. While this represents background signal in Hi-C, it indicates loss of mitochondrial integrity and mitochondrial DNA leakage into the cytoplasm, as has been observed in earlier studies (Yu et al., 2020).

The spatial conformation of the *C9orf72* locus, and its flanking domains do not appear to be affected by the hexanucleotide repeat expansion: while the locus displays differences in conformation, compartmentalization, and boundary positions as cells reprogram, differentiate and mature, these changes are similar in ALS cases and controls. Due to the fact that the ALS-derived cells carry both an expanded allele and a normal allele, the genomic data represents a mixture of both. This may lower our power to detect more subtle effects.

Finally, we find variation in severity of chromosome conformation defects in the four clones of motor neurons derived from ALS patients, even for clones derived from the same patient. We observed some differences during reprogramming as well, that correlate with the severity of the defects observed in the motor neurons. This suggests that both the process of reprogramming and maturation has some variation unrelated to the genotype of the donor. It would be interesting to identify the factors that influence the severity of the ALS defect in motor neurons.

In summary, we describe dynamic changes in chromosome organization during neuronal maturation and identify defects in this process in ALS neurons. Future studies can reveal whether these defects themselves contribute to motor neuron defects or whether they represent another downstream hallmark of the disease.

## Code Availability

No new code was generated for this study. Hi-C mapping pipeline distiller-nf is available on Github: https://github.com/mirnylab/distiller-nf. Downstream analysis tools pairtools and cooltools are available through https://github.com/mirnylab/pairtools and https://github.com/mirnylab/cooltools. Code used for analysis of ATAC-seq data can be found at Github: https://github.com/dekkerlab/Ozgun-ALS.

## Acknowledgements

We thank Dr. Jemeen Sreedharan for his valuable scientific input during the initiation of this study. We sincerely thank the Flow Cytometry Core Facility at UMass Chan Medical School for their expert support and assistance in sorting motor neurons for this study. JD is an investigator of the Howard Hughes Medical Institute. Dr. Brown gratefully acknowledges support for his laboratory from ALS One, The Angel Fund for ALS Research, NIH awards from the NINDS (R21NS139270, RO1NS111990, R01NS104022) and NIGMS (R01GM1502730) and also from the DOD (HT9425-24-1-0830 AL230185).

## Author contributions

J.D., R.H.B., and Ö.U. conceived and designed the study. K.E. and J.R.K. supervised the reprogramming and differentiation experiments. Ö.U., M.O., N.W., J.R.K., H.B., and Z.S.A.U. performed the experiments. J.D., R.H.B., Ö.U., S.S., M.O., A.S., Ö.A., and B.A.-O. analyzed the data. J.D., R.H.B., and Ö.U. wrote the manuscript.

## DECLARATION OF INTERESTS

Job Dekker is a member of the scientific advisory board of Arima Genomics, San Diego, CA, USA and Omega Therapeutic, Cambridge, MA, USA. Elsevier is the current employer of Zeynep Sena Agim Uslu. Uslu confirms that research was completed before this author’s employment at Elsevier and that the peer review was fully independent of this person.

## SUPPLEMENTAL INFORMATION

### Materials and Methods

#### Primary cell lines and cell reprogramming

Two primary normal human dermal fibroblast cells (control-1 and control-2), one male and one female individual were obtained from Coriell institute. ALS patients with C9orf72 HRE primary fibroblast lines were obtained from the repository of the Brown lab (C9-ALS-1 and C9-ALS-2). All primary fibroblasts (PF) were cultured in DMEM, high glucose, GlutaMAXTM supplement with pyruvate (Gibco, cat. no. 10569016) with 10% fetal bovine serum (Sigma, cat. no. F2442) at 37°C in 5% CO2. Cells were passaged every 3-4 days before reaching confluency. For PF Hi-C, cells were harvested at day 4 and day 16. All PF lines were reprogrammed using SeV-based reprogramming kit (CytoTune 2.0 sendai reprogramming kit, life technologies). Protocol is applied as indicated in manufacturer’s instructions. Briefly, 1-2.5 x 105 fibroblast cells were seeded in three wells of a 6-well plate and cultured 2 days to reach 50–65% confluency. SeV reprogramming vectors (CytoTune 2.0, KOS; CytoTune 2.0 hL-Myc; CytoTune 2.0 hKlf4) were added as guided in the protocol. Spent media was replenished every other day until day 7. On day 7 after viral exposure, cells were detached using Trypsin-EDTA (0.25%) (Gibco, cat. no. 25200056), and re-plated on 6-well plates coated with Matrigel (Corning, cat. No. 354277) and maintained in mTeSR1 medium (Stem Cell Technologies) with daily feeding. After 4-5 weeks post-transduction, human iPSC colonies became visible and expanded to be separated from undifferentiated fibroblasts. For each primary fibroblast line that is derived, two IPSC clones were maintained and sent to karyotype analysis for proper chromosome integrity (WiCell).

#### Maintenance of IPSCs and differentiation of IPSCs into motor neurons

IPSCs were maintained in mTeSR1 medium (Stem Cell Technologies) on 6-well culture plates coated with Matrigel (Corning, cat. no. 354277) at 5% CO2 and 37 °C incubators. IPSCs were incubated in DPBS (Gibco, cat. no. 14190144) containing 1 mM EDTA for few minutes and then dissociated with pipette up and down. Small aggregates of cells with very low density were passaged into new matrigel coated plates containing mTeSR1 medium with 10 μM ROCK inhibitor (Sigma, Y-27632). Fresh mTeSR1 medium was added 24 hours later to remove ROCK inhibitor from the cells. Cells were harvested when they reach 80-90% confluency for Hi-C and RNA-seq experiments. Motor neuron differentiation was performed as described previously (Klim et al., 2019). In brief, IPSCs were plated on Matrigel-coated culture plates with mTeSR1 medium containing 10 μM ROCK inhibitor. When cells reached 100% confluency, media was changed to differentiation medium (%50 Neurobasal (Gibco, cat. no. 21103049) and 50% DMEM-F12 (Gibco, cat. no. 11320033) containing 1x B-27 supplement (Gibco, cat. no. 17504044), 1x N-2 supplement (Gibco, cat. no. 17502048), 1x GlutaMAX supplement (Gibco, cat. no. 35050079) and 1x MEM Non-Essential Amino Acids (NEAA) Solution (Gibco, cat. no. 11140076)). The following treatments with small molecules were carried out for days 1-6: 10 μM SB-431542 (sigma, cat. no. S4317), 100 nM LDN-193189 (Stemgent/reprocell, cat. no. 04-0074-02), 1 μM retinoic acid (RA) (Sigma, cat. no. R2625) and 1 μM Smoothened agonist (SAG) (Sigma-aldrich, cat. no. 566660); for days 7-14, 5 μM DAPT (Sigma-aldrich, cat. no. D5942), 4 μM SU-5402 (Sigma-aldrich, cat. no. SML0443), 1 μM RA and 1 μM SAG.

Fluorescence-activated cell sorting (FACS) of immature motor neurons At day 14 after neural differentiation, cultures were dissociated to single cells using StemPro accutase treatment (Gibco, cat. no. A1110501) containing DNAseI (Worthington biochem, cat. no. LK003172) for 30-45 minutes at 37 °C. Gentle pipetting with a 1,000 μl pipetman was used (10–20 up and downs) for complete cell dissociation. Cells were passed through a 40 μm filter and spun down, washed ×1 with DPBS and resuspended in sorting buffer (1x cation-free PBS (Gibco, cat. no. 14190144), 15 mM HEPES (Gibco, cat. no. 15630080), 1% BSA (Sigma, cat. no. A8412-100ML), 1x penicillin-streptomycin (Gibco, cat. no. 15140122), 1 mM EDTA (Invitrogen, cat. no. AM9260G). Single cell suspensions were incubated in sorting buffer containing NCAM (BD Bioscience, cat. no. BDB557919; 1:200 dilution) and EpCAM (BD Bioscience, cat. no. BDB347198; 1:50 dilution) antibodies for 30 minutes and washed with sorting buffer and then resuspended in sorting buffer with DAPI (Invitrogen, cat. no. D3571). The sorted EpCAM-negative NCAM-positive cells were collected and plated onto Poly-L-lysine coated plates (Sigma, P5899) with motor neuron media (Neurobasal medium, 1x N-2 supplement, 1x B-27 supplement, 1x GlutaMax and 1x MEM NEAA solution) with 10 μM ROCK inhibitor and 10 ng per ml of the following neurotrophic factors: glial cell-derived neurotrophic factor (GDNF) (R&D systems, cat. no. 212-GD-010/CF), brain-derived neurotrophic factor (BDNF) (R&D systems, cat. no. 11166-BD-010) and ciliary neurotrophic factor (CNTF) (R&D systems, cat. no. 257-NT-010/CF). Half of the spent media was replenished every 2-3 days. Motor neurons were cultured up to 6 weeks until they were harvested for Hi-C, RNAseq and ATAC-seq.

#### RNA Fluorescence in situ Hybridization (FISH)

RNA FISH was performed using a 5’ end Cy3-conjugated (G2C4)4 DNA probe to detect sense RNA of the C9orf72 gene. Fibroblasts were fixed in 1x DEPC-treated PBS containing 4% PFA for 15 mins. Fixed cells were washed three times with PBS and were permeabilized with 0.5% triton X-100 (Sigma, cat. no. x100-100ml) in PBS for 10 mins. Then cells were incubated at 56°C in hybridization buffer including 40% formamide (Fisher, cat. no. BP227-500), 2x saline sodium citrate (SSC) buffer, 0.1% Tween-20 and Herring sperm DNA (Promega, cat. no. D1815) for 1 hour. After that cells were incubated with hybridization buffer containing Cy3-conjugated (G2C4)4 DNA probe. Cells were washed twice in pre-warmed wash buffer (40% formamide, 2x SSC, 0.1% Tween-20 (ThermoFisher, cat. no. 85113)) and then in stringency wash buffer (0.2x SSC and 0.1% Tween-20) at 55°C. Samples were then mounted in Prolong Diamond Antifade reagent with DAPI (ThermoFisher, cat. no. P36962) for at least 24 hours.

#### Southern Blot

8-10 μg genomic DNA (gDNA) was extracted for each sample using gentra puregene blood kit (Qiagen). The gDNA was digested with AluI (NEB, cat. no. R0137L) and DdeI (NEB, cat. no. R0175L) restriction enzymes at 37°C overnight. DNA samples were loaded on 0.8% agarose gel and separated by electrophoresis for 2 hours. Then, the trimmed gel was washed with depurination Buffer (0.25M HCl) for 20 min and rinsed with ddH O briefly three times. The gel was shaken in denaturation buffer (0.5M NaOH/ 1.5M NaCl) for 40 min and rinsed again with ddH O briefly three times. Finally, the gel was incubated in neutralization buffer (0.5M Tris-HCl/ 1.5M NaCl; pH 7.5) for 30 min and rinsed briefly with ddH O three times. Then, samples were transferred to a positively charged nylon membrane (Roche) using conventional southern blot sandwich method. Transferred gDNA on the nylon membrane was cross-linked by UV (1200J) and incubated with Easy HYB buffer (Roche-Cat no. 11603558001) containing 150μl of Herring Sperm DNA (Promega, cat. no. D1815) (Blocking buffer). The blot was hybridized with a digoxigenin-labeled G2C4 DNA probe in fresh Easy HYB buffer overnight at 55°C. Next day, the blot was firstly washed twice with low stringency wash buffer (2xSSC + 0.1% SDS) at room temperature for 15 min and then washed twice with high stringency wash buffer (0.1x SSC + 0.1% SDS) at 68°C for 30 min. Finally, blot was prepared for detection using DIG wash and block buffer kit (Roche, cat. no. 11585762001). The digoxigenin-labeled probe was detected with anti-digoxigenin antibody and CDP-Star reagent (Roche, cat. no. 12041677001).

#### Immunoblot analysis

Motor Neuron week 3 and 6 total proteins were extracted using RIPA buffer containing complete mini protease inhibitor cocktail tablets (Roche, cat no. 11836153001). Protein concentration was calculated using Pierce BCA protein assay kit (ThermoFisher, cat no. 23225). 5µg of total protein was loaded onto either 12% or 4-12% Tris-Glycine gels (Invitrogen). Proteins were separated using a mini gel tank (Invitrogen), in 1x TG-SDS running buffer (Invitrogen) at 120 volts for 1.5 hours. Proteins were transferred to nitrocellulose membranes using an iBlot 2 device (Invitrogen) following manufacturers protocols. Membranes were immediately placed into Intercept blocking buffer (LICOR) and incubated at RT for 1 hour. After blocking, the blots were placed into primary antibody solution which consisted of Intercept blocking buffer (LICOR), 0.1% Tween-20 (Fisher), and various combinations of the following primary antibodies; rabbit anti-RAD21 (Abcam, cat no. ab992, 1:1000), rabbit anti-H3 (Abcam, cat no. ab1791, 1:1000), goat anti-βactin (Abcam, cat no. ab8229, 1:3000), rabbit anti-HP1α (Abcam, cat no. ab109028, 1:1000), Rabbit anti-TDP-43 (Proteintech, cat no. 10782-2-ap, 1:1000). Blots were incubated with primary antibody solution overnight at 4°C. The next day, blots were washed 4x with PBS-T, 5 minutes for each wash. The blots were then transferred to secondary antibody solution; intercept blocking buffer (LICOR), 0.1% tween-20 (fisher), and the appropriate Infrared tagged secondary antibodies (LICOR). Blots were incubated in light blocking containers for 1 hour at RT before being washed 4x with PBS-T (5 minutes each), followed by one final wash in PBS for 5 minutes. Blots are stored in PBS in light blocking containers until being visualized on an Odyssey Infrared imager. Densitometry was determined using the Odyssey V3.0 software.

#### RNA-seq library preparation and data processing

Total RNAs of fibroblast, IPSC and MN lines were extracted using Direct-zol RNA Microprep Kit (Zymo, R2061) with DNAseI digestion. RNA-seq libraries were prepared using KAPA RNA HyperPrep with RiboErase as instructed in manufacturer’s protocol (Roche, cat. no. 08098131702). RNA-seq libraries were amplified by 8 PCR cycles using Illumina adapters and primers (TruSeq DNA LT kit Set A 15041757). The libraries were sequenced using 150bp paired end reads on Illumina HiSeq4000.

Fastq reads of RNA-seq libraries were processed using DolphinNext/ViaFoundry platform (Yukselen et al., 2020). Briefly, adapter removal was performed to remove adapter containing sequences from the fastq reads and then sequential mapping was run to remove rRNAs, miRNAs, snRNAs, piRNAs and tRNAs. RSEM was performed using STAR to align reads to reference transcriptome (Gencode v43) to estimate gene and isoform expression levels. To identify differentially expressed genes, adapter removal and low-quality base trimming were performed with Cutadapt. Gene expression was quantified using Salmon (Patro et al., 2017).

DESeq2 (Love et al., 2014) pipeline was run using UMassMed dolphinnext/viafoundry servers (dolphinnext.umassmed.edu/viafoundry.umassmed.edu). TPM (transcript per million) count tables were used in gene expression comparisons among cell-types.

For significant differentially expressed gene selection, the following cut-offs were used: Genes that have less than 100 counts among the samples were removed for the DESeq2 analysis. Furthermore, >1.5 fold-change and p-values<0.05 were applied. For gene set enrichment analysis (GSEA), upregulated and downregulated gene lists were analyzed using Enrichr (Xie et al., 2021), GO Biological Process 2025.

#### Omni ATAC-seq and analysis

OmniATAC-seq was performed as published previously (Corces et al., 2017). Briefly, motor neurons were washed and lysed while adherent to the plate using 0.1% NP-40, 0.1% Tween-20, 0.01% digitonin, 10 mM Tris-HCl (pH 7.4), 10 mM NaCl and 3 mM MgCl2. Chromatin from approximately 50,000 nuclei was transposed using 10uL Tn5 from the Nextera DNA library prep kit (Illumina, cat. no. FC-121-1030) for 30 min at 37C in the presence of 0.01% digitonin and 0.1% Tween-20. DNA was immediately purified using Qiagen MinElute Kit (Qiagen, cat. no. 28004) and kept at-20°C until PCR amplification. Cycle number for the amplification of the DNA was determined using qPCR. All samples were amplified using less than 8 PCR cycles. Primers were removed from amplified libraries using AMpure XP beads (Beckman Coulter, cat. no. A63881). The libraries were sequenced on illumina HiSeq4000 for 50bp paired end reads.

ATAC-seq data were mapped and processed as described previously (Oomen et al., 2019). Briefly, reads were trimmed from 24 bp and aligned to hg38 using Bowtie2 with maximum mapping length of 2000 bp (Langmead and Salzberg, 2012). Mapped reads were filtered for mapping quality, mitochondrial reads, PCR duplicates and blacklisted. For downstream analysis, each end +5 bp was taken and analysis was continued treating the reads as single end reads.

Further downstream processing was performed using BEDtools (Quinlan and Hall, 2010) and HOMER (Heinz et al., 2010).

#### Footprinting analysis of ATAC-seq data

We used MACS3 (Zhang et al., 2008) to find ATAC-seq peaks of accessibility using default parameters with a shift/extend of-75/+150. The union peak set was merged using bioframe (Open2C et al., 2024). Footprinting analysis was performed using TOBIAS (Bentsen et al., 2020) across the union peak set generated for the “all control MN, all ALS MN, MN-w0, MN-w3, and MN-w6” conditions. First, ATACseq signal was corrected for Tn5 insertion bias using the filtered alignments (ATACorrect). The corrected signal was then used to calculate a continuous footprinting score (ScoreBigwig), determining local regions of decreased accessibility. Finally, mean transcription factor footprinting scores as well as pairwise differential “binding” scores (BINDetect) were determined for 841 conserved vertebrate motifs (Fornes et al., 2020), across the union peak set. From BINDetect scores, bound change values of transcription factors with significant p-values for MN-w6 vs MN-w0 comparison were extracted as a list. Pairwise bound change scores of the transcription factors listed for MN-w0 vs MN-w3, MN-w3 vs MN-w6, ALS vs control, MN-w0 vs ALS, MN-w3 vs ALS were also extracted and comparisons were plotted as a heatmap. Additionally, log2foldchange values for available genes in the list were also plotted as a heatmap for comparison.

#### Hi-C library preparation

Hi-C 2.0 was performed as described previously (Belaghzal et al., 2017). 5-10 million cells for each line were fixed with 1% formaldehyde in PBS for exactly 10 minutes at room temperature. 125mM glycine was added and incubated to quench the formaldehyde and terminate the cross-linking reaction for 5 mins at room temperature (RT) and then put on ice for 15 minutes. Cells were scraped from the plates, spun. Supernatant was discarded and then proceeded to cell lysis. 1 ml of ice-cold lysis buffer (10mM Tris-HCl pH8.0, 10mM NaCl, 0.2% Igepal CA630 (Sigma, cat. no. I8896-50ML), 10μL of 10X protease inhibitors (Thermofisher, cat no. 78438)) was added on the fixed cells and put on ice for 15 minutes. Then, cells were lysed with a dounce homogenizer and pestle A (Kimble Kontes cat. no. 885303-0002) by moving up and down 30 times, incubating on ice for one minute followed by 30 more strokes with the pestle. For motor neuron preps, this step was repeated one more time. The suspension was centrifuged for 5 minutes at 2,000g at RT and supernatant was discarded, and the pellets were washed twice with ice-cold 500μL of 1x NEBuffer 3.1. Then the pellet was resuspended in 720μL of 1x NEBuffer 3.1. 38μL of 1% SDS was added to each tube and incubated at 65°C for 10 minutes to solubilize chromatin and then put on ice to cool down. 43μL of 10% Triton X-100 was added to quench SDS. Next, chromatin was digested by adding 400 Units DpnII enzyme (NEB) at 37°C for 16-18 hours digestion within a shaker and then incubated at 65°C for 20 minutes to inactivate the enzyme and then directly put on ice. Biotin fill-in was performed to mark the ends of digested DNA fragments. 60μL of biotin fill-in master mix (1X NEB 3.1, 0.25mM dCTP, 0.25mM dGTP, 0.25mM dTTP, 0.25mM biotin-dATP (ThermoFisher.cat. no. 19524016), DNAvPolymerase I, large Klenow fragment (NEB, cat. no. M0210)) was added to each tube and incubated at 23°C for 4 hours. Biotin filled ends were ligated by adding 665μL of ligation mix (243μL ultra-pure water (Invitrogen, cat. no. 10977015), 240μL of 5X ligation buffer (1.8X) (Invitrogen, cat. no. 46300018), 120μL 10% Triton X-100, 12μL of 10mg/mL BSA and 50μL T4 DNA ligase (Invitrogen, cat. no. 15224090)) and incubated at 16°C for 4 hours. Then, crosslink was reversed by adding 50μL of 10mg/mL proteinase K (Fisher scientific, cat. no. BP1750I-400) and incubated at 65°C for 3 hours. Another 50μL of proteinase K was added and incubated overnight at 65°C. DNA was extracted by adding 1:1 volume of saturated phenol pH 8.0: chloroform (25:24) (Fisher scientific, cat. no. AC327115000), and vortexed for 15 seconds and then transferred to phase-lock tubes and spun at 16,000g for 5 minutes to separated inorganic and organic phases. Ligated DNA was precipitated by adding a volume of 3M sodium acetate pH 5.2 equal to 10% of the initial volume and 2 volumes of ice-cold ∼100% ethanol (Fisher scientific, cat. no. BP2818500). Tubes were mixed well and incubated for at least two hours at-80°C. Next, they were centrifuged at 16,000g at 4°C for 30 minutes. The pellet was dissolved in 500μL of 1X Tris-EDTA (TE) (IDT, cat. no. 11-05-01-09) and transferred to a 0.5 ml amicon ultra centrifuge filter (EMD Millipore, cat. no. UFC5030BK). Another 500μL of 1X TE was added to the samples, then, columns were spun at max g for 5 minutes and the flowthrough was discarded. This step was repeated four times. The pellet was dissolved in 50μL of ultra-pure water. 1μL of 10 mg/mL RNAase A was added and incubated at 37°C for 30 minutes. Finally, DNA was quantified by loading on a 1% agarose gel. Hi-C libraries were treated with T4 DNA polymerase to remove unbiotinylated DNA fragment ends. Up to 5μg DNA of Hi-C library was added to a 50μL reaction tube containing 5μL of 10X NEBuffer 3.1, 0.025mM dATP, 0.025mM dGTP and 15U T4 DNA polymerase (NEB, cat. no. M0203L). Each tube was incubated at 20°C for 4 hours, and then inactivated at 75°C for 20 minutes. Appropriate volume of 1X TE was added to make it 130μL. DNA fragments were sheared to a size of ∼200bp using a covaris instrument (Duty Factor 20%, Cycles per Burst 200, peak power 50, average power 17.5 and process time 180 sec). Size selection was applied to enrich DNA fragments between 100-300bp using ampure XP fractionation (Beckman Coulter, cat. no. A63881) and the pellet was eluted with 50μL of ultra-pure water. The quality of DNA fragments was checked on a 2% agarose gel. Next, end repair was performed using 45μL of Hi-C samples transferred to each PCR tube, and 25μL of the end-repair mix containing 3.5X NEB ligation buffer (NEB, cat. no. B0202S), 17.5mM dNTP mix, 7.5U T4 DNA polymerase (NEB, cat. no. M0203L), 25U T4 polynucleotide kinase, 2.5U Klenow polymerase Polymerase I, was added and incubated at 37°C for 30 minutes. Then tubes were additionally incubated at 75°C for 20 minutes to inactivate the enzymes. 50μL of MyOne streptavidin C1 bead mix (ThermoFisher, cat. no. 65001) was transferred to a 1.5mL tube. The beads were washed twice by adding 400μL of TWB (5mM Tris-HCl pH=8, 0.5mM EDTA, 1M NaCl, 0.05% Tween20) and incubated for 3 minutes at RT. Then, beads were resuspended in 400μL of 2X binding buffer (10mM Tris-HCl pH=8, 1mM EDTA, 2M NaCl) and mixed with 400μL of DNA fragments from the previous step. The mixture was incubated for 15 minutes at RT with rotation. Beads with bound DNA were washed with 400μL of 1X binding buffer and then washed once with 100μL of 1X NEB2.1. Finally, beads containing DNA were resuspended in 41μL of 1x NEB2.1. 9μL of A-tailing mix (5μL of NEB buffer 2.1, 5μL of 1mM dATP, 3U Klenow exo (NEB, cat. no. M0212S)) was added on top of 41μL of beads. The reaction was incubated at 37°C for 30 minutes, then at 65°C for 20 minutes, and then cooled to 4°C. The beads containing bound DNA were washed twice with 100μL of 1X T4 DNA ligase buffer and then resuspended in 35μL 1X T4 DNA ligase buffer. For final step of library preparation, PCR amplifications, TruSeq DNA LT kit Set A (Illumina, cat. no. 15041757) was used. Three different PCR cycles (6-8-10) were tested to pick best amplification cycle for each sample. 6-8 PCR cycles was performed for final library preparation using primer mix and Master Mix of TruSeq DNA LT kit Set A. Final library products were cleaned up using Ampure XP beads and sequenced using 50bp paired end reads on an Illumina HiSeq4000 instrument.

#### Hi-C data processing

Distiller (https://github.com/mirnylab/distiller-nf) pipeline was used to process Hi-C fastq files. First, fastq reads were mapped to hg38 using bwa mem. Second, mapped reads were parsed and classified using the pairtools package (https://github.com/mirnylab/pairtools) to get pairs files. PCR duplicates were removed, and pairs were filtered using mapping quality scores (MAPQ > 30). Data were binned into 1, 2, 5, 10, 25, 50, 100, 250, 500 and 1000kb resolutions. Finally multiresolution cooler files were created using the cooler package (https://github.com/mirnylab/cooler.git). Contact matrices were normalized using the iterative correction procedure published previously (Imakaev et al., 2012).

#### Cis and Trans Ratio

Cis and trans percentages were calculated by dividing the total interactions between chromosomes with the sum of interactions within and between chromosomes. The numbers of interactions captured within and between chromosomes were obtained from pairtools.

#### Compartment Analysis

A and B compartments were assigned using an eigenvector decomposition procedure implemented in the cooltools package. Eigenvector decomposition was performed on observed-over-expected cis contact matrices at 25 kb and 100kb bin size for every chromosome. The first eigenvectors (EV1/Eig1) positively correlated with the gene density were used to call A or B compartment identity for each bin. Compartmentalization strengths were calculated using saddle-plots as implemented in cooltools version 0.5.4 (Imakaev et al., 2012). Briefly, saddle plots were generated by sorting the eigenvector values from lowest to highest (B to A). Then, sorted maps were normalized with their expected interaction frequencies. The upper left corner of the interaction matrix represents the strongest B-B interactions and the lower right for strongest A-A interactions. Upper right and lower left are for B-A and A-B respectively. To quantify and compare saddle plots of different samples, the strongest 20% of BB and strongest 20% of AA interactions were taken and divided them by the sum of AB and BA (AA)/(AB+BA) and (BB)/(AB+BA).

For generating centromere-telomere plots (Figure 2H and 7D), the saddles were piled up using generated centromere and telomere CT track. Centromere-telomere tracks were generated for every chromosome at 100kb bin size, by assigning values from 0 to 1 for p arm, and 1 to 2 for q arm with 1 representing centromeric regions, and 0 and 2 the telomeric regions. Cooltools-saddle function of trans pile was used to generate the pileups for chromosomes 1 to 22.

For arm specific telomeric interaction plots (Figure 5h and 7E), telomeric interactions were obtained from 250kb matrices. First four 250kb bins of each telomeric end with valid balance weight score (1 Mb) was used to obtain total trans and cis telomeric interactions from balanced cooler matrices for all autosomal chromosomes. First diagonal scores indicate cis telomeric end interaction value for each chromosome. Other squares with 4 sub squares indicate trans telomeric arms’ interaction scores between chromosome pairs.

#### Insulation score analysis

insulation scores were calculated using cooltools (https://github.com/open2c/cooltools/blob/master/cooltools/cli/diamond_insulation.py) as implemented from Crane et al (Crane et al., 2015). The insulation analyses were done on 10kb binned data with a 100kb window size. First two diagonals were excluded for insulation score calculations. Strong boundaries were defined by the boundary strength values greater than or equal to 0.1 for comparison of week 6 and week 0 samples. Genomic bin that has “nan” values in at least 1 bin out of 5 flanking right or left bins were removed. The regions that are common and have an overlap of 5kb were considered as intersected regions. For generating stack-ups, bbi.stackup() function was used, with 100kb flanking regions. The compartments were called at 25kb resolution and insulation at 10kb with a 100kb sliding window. The regions were arranged based on compartment change status across the middle bin with regions sorted in descending order of EV value as a second key of sorting. The insulation stack-ups were normalized by subtracting the mean of first three and last three bins of each row from itself.

## Supplementary figures

**Figure S1 (associated with Figure 1). Generating induced pluripotent stem cells from primary fibroblasts**

a. Schematic view of the study design.

b. Schematic view of IPSC generation from primary fibroblast using CytoTune iPS 2.0 Sendai Reprogramming Kit.

c. Formation of Embryoid bodies (EB) to generate three germ lines from IPSC clones. TUBB3 (Tuj1): βIII-tubulin (Neural, ectodermal); SMA: Smooth muscle Actin, mesodermal; SOX17: SRY (sex determining region Y)-box 17 (Vascular, endodermal).

**Figure S2 (associated with Figure 2). Chromatin architecture of cell type–specific genes in PFs, iPSCs, and MNs.**

Hi-C contact matrices at 25 kb resolution with A/B compartment tracks (first eigenvector, EV1) are shown for PFs, iPSCs, and MNs (columns). Each heatmap is centered on the gene of interest, with ±2 Mb of flanking regions. (a) COL1A2 (PF), SOX2 (iPSC), MNX1 and MAP2 (MN); (b)

ANXA2 (PF), ESRG (iPSC), NRG1 and FOXP1 (MN); (c) COL1A1 (PF), L1TD1 (iPSC), NEFH and PRPH (MN); (d) GFAP, S100B, TMEM119, and AIF1 (glial markers). Top panels display GENCODE gene annotations for each region.

**Figure S6 (associated with** Figure 6**). Analysis of TAD boundaries during maturation.** Boundary strength distribution of insulation scores sorted by log 10 of their boundary strengths in EP-W0 (Total:48366 Strong:17148), MN-W0 (Total:47012 Strong:16211), MN-W3 (Total:36689 Strong:14521) and MN-W6 (Total:33008 Strong:13599) (strong boundary > log10 - 1) Insulation scores and boundary strength are calculated at 10 kb resolution with 100 kb window size.

**Figure S7 (associated with Figure 7). Characterization of the *C9orf72* HRE mutation in ALS patients and chromatin architecture in C9-ALS PFs, iPSCs, and MN cells compared to control cells.**

a. RNA-fluorescence in situ hybridization (RNA-FISH) in primary fibroblasts of control-1, C9-ALS-1 and C9-ALS-2 individuals using a Cy3 labeled (G2C4)_4_ DNA probe. White arrows indicate (G4C2)_n_ containing RNA foci. Blue: DAPI; Red: Cy3-conjugated (G2C4)_4_ DNA probe.

b. Southern blot analysis of fibroblasts from control-1 and C9-ALS-1. Repeat-containing fragments were detected using a region-specific, 5′ DIG-labeled PCR-amplified probe. The expanded allele shows a repeat size of approximately 800–1000.

c. Southern Blot analysis of IPSCs and fibroblasts obtained from C9-ALS-1 and C9-ALS-2 patients. Repeat expansions were detected using a 5’DIG-(G4C2)_5_-DIG-3’ DNA probe revealing a ∼5-6 kb band corresponding to an expanded repeat size of 800–1000. Positive control: ALS patient with a known *C9orf72* HRE.

d. Hi-C contact matrices of chromosome 12 interactions with A/B compartment calls (100k bin size) in PFs, two IPSCs and two MNs from control-1, C9-ALS-1 and C9-ALS-2, individuals (Eig1= Eigenvalue 1).

e. Zoomed-in Hi-C contact matrices showing compartmental and TAD interactions in the 45–65 Mb region of chromosome 12 (100 kb bin size) for PFs, two iPSCs, and two MNs from control-1, C9-ALS-1, and C9-ALS-2 individuals (Eig1 = first eigenvector).

Quantification of compartment strengths (AA and BB) using cis-interactions derived from saddle plots of PFs, iPSCs, and MNs from C9-ALS-1 and C9-ALS-2.

g. Genome-wide cis and trans saddle plots of chromatin interactions in PFs, IPSCs and MNs from control-1, control-2, C9-ALS-1 and C9-ALS-2 individuals.

h. Hi-C contact matrices (25 kb bin size) focused on the 26–30 Mb region of chromosome 9 in PFs, two iPSCs, and two MNs from control-1, control-2, C9-ALS-1, and C9-ALS-2. Bottom panels display GENCODE gene annotations for each region.

i. Venn diagram showing the overlap of strong chromatin boundaries between week 0 and week 6 as derived from Figure 6G. Boundaries include W0-specific (n = 7109), shared (W0 & W6; n = 7073), and W6-specific (n = 5115) sets.

j. Aggregated insulation score pileups of strong boundaries (union from panel i) across PFs, iPSCs, and MNs from control-1, control-2, C9-ALS-1, and C9-ALS-2 individuals.

k. Stack-up plots of insulation scores detected at the strong boundaries with flanking 100kb from each side for control-1, control-2 and C9-ALS-1 and C9-ALS-2 individuals. W0-specific, intersection, and W6-specific strong boundaries detected in figure S6I are used.

**Figure S8 (associated with Figure 8). Transcriptome and genome accessibility impacts of C9orf72 HRE Mutation in ALS Motor Neurons**

a. Expression levels of genes associated with the synapse organization biological process were plotted across four conditions: cntrl-W0, cntrl-W3, cntrl-W6, and ALS-W6. The panels represent three comparisons. Left panel: up. genes in cntrl-W6 vs cntrl-W0 (# of Genes: 53); mid panel: down. genes in cntrl-W6 vs cntrl-W3 (# of Genes: 17); right panel: down. genes in ALS-W6 vs cntrl-W3 (# of Genes: 32).

b. Expression levels of genes associated with the axonogenesis biological process were plotted across four conditions: cntrl-W0, cntrl-W3, cntrl-W6, and ALS-W6. The panels represent three comparisons. Left panel: up. genes in cntrl-W6 vs cntrl-W0 (# of Genes: 69); mid panel: down. genes in cntrl-W6 vs cntrl-W3 (# of Genes: 27); right panel: down. genes in ALS-W6 vs cntrl-W3 (# of Genes: 39).

c. Expression levels of genes associated with the Anterograde trans-synaptic signaling biological process were plotted across four conditions: cntrl-W0, cntrl-W3, cntrl-W6, and ALS-W6. The panels represent three comparisons. Left panel: up. genes in cntrl-W6 vs cntrl-W0 (# of Genes: 64); mid panel: down. genes in cntrl-W6 vs cntrl-W3 (# of Genes: 18); right panel: down. genes in ALS-W6 vs cntrl-W3 (# of Genes: 30).

d. Expression levels of genes associated with the cellular respiration biological process were plotted across four conditions: cntrl-W0, cntrl-W3, cntrl-W6, and ALS-W6. The panels represent two comparisons. Left panel: up. genes in cntrl-W6 vs cntrl-W0 (# of Genes: 38); right panel: down. genes in ALS-W6 vs cntrl-W3 (# of Genes: 16)

e. Expression levels of genes associated with the proton motive force-driven ATP synthesis biological process were plotted across four conditions: cntrl-W0, cntrl-W3, cntrl-W6, and ALS-W6. The panels represent two comparisons. Left panel: up. genes in cntrl-W6 vs cntrl-W0 (# of Genes: 25); right panel: down. genes in ALS-W6 vs cntrl-W3 (# of Genes: 10)

f. Expression levels of genes associated with the vacuolar/lysosomal lumen acidification biological process were plotted across four conditions: cntrl-W0, cntrl-W3, cntrl-W6, and ALS-W6. The panels represent two comparisons. Left panel: up. genes in cntrl-W6 vs cntrl-W0 (# of Genes: 15); right panel: down. genes in ALS-W6 vs cntrl-W3 (# of Genes: 1)

g. Heatmap displaying the differential binding scores of transcription factors (TFs) with the most significantly upregulated and downregulated binding activity. Genome-wide TF binding dynamics were analyzed using footprinting analysis of ATAC-seq data from control MNs (W0, W3, and W6) and C9-ALS MNs (W6). Differential binding scores for each TF were calculated using TOBIAS.

## Notes

### Summary of Updates

The order of the authors has been corrected.

