## Supplementary figures and images for "Dynamic changes in chromosome and nuclear architecture during maturation of normal and ALS C9orf72 motor neurons"

### Supplementary Figure S1

#### Supp. Figure 1

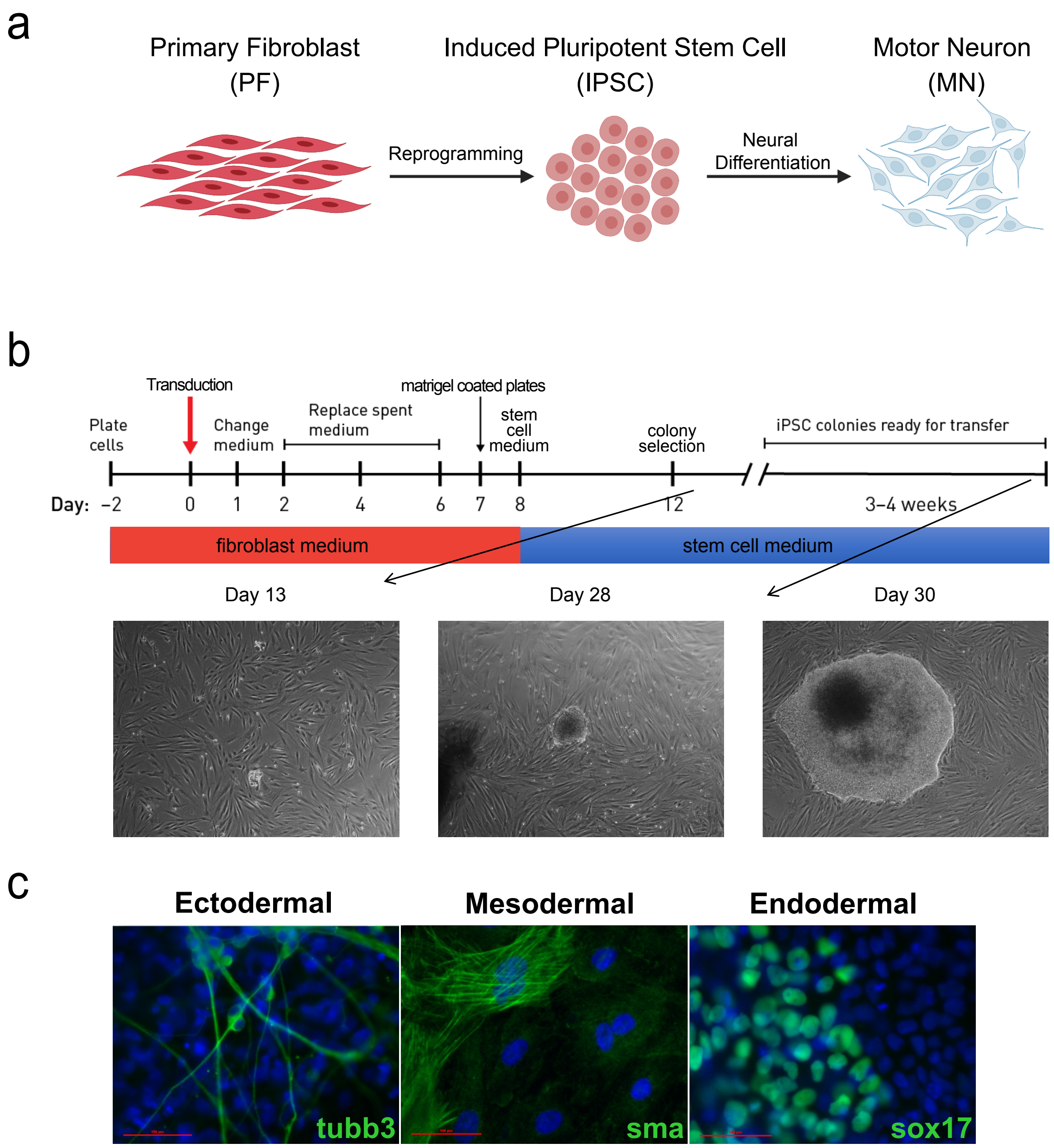

### Supplementary Figure S2AB

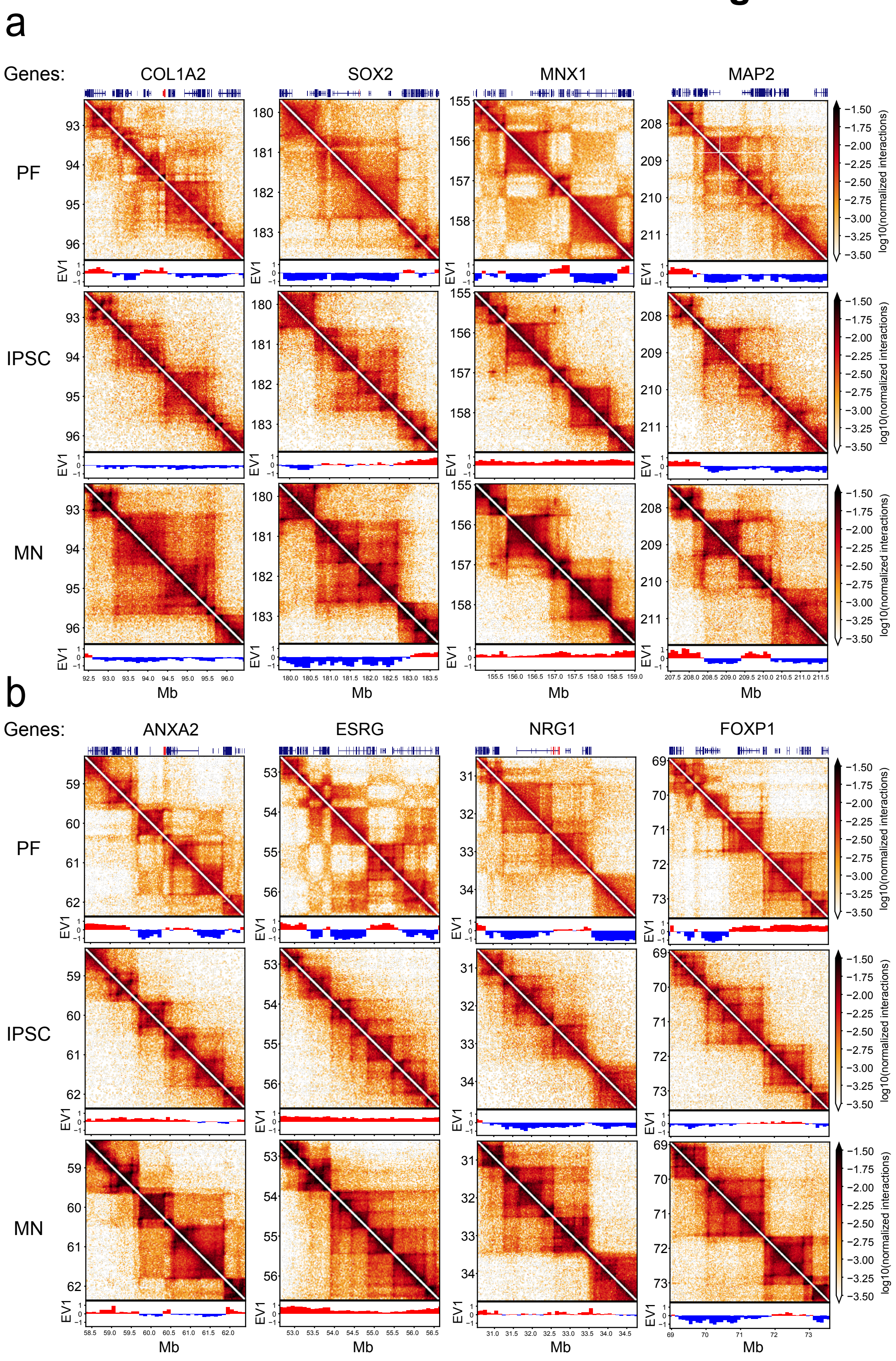

### Supplementary Figure S2CD

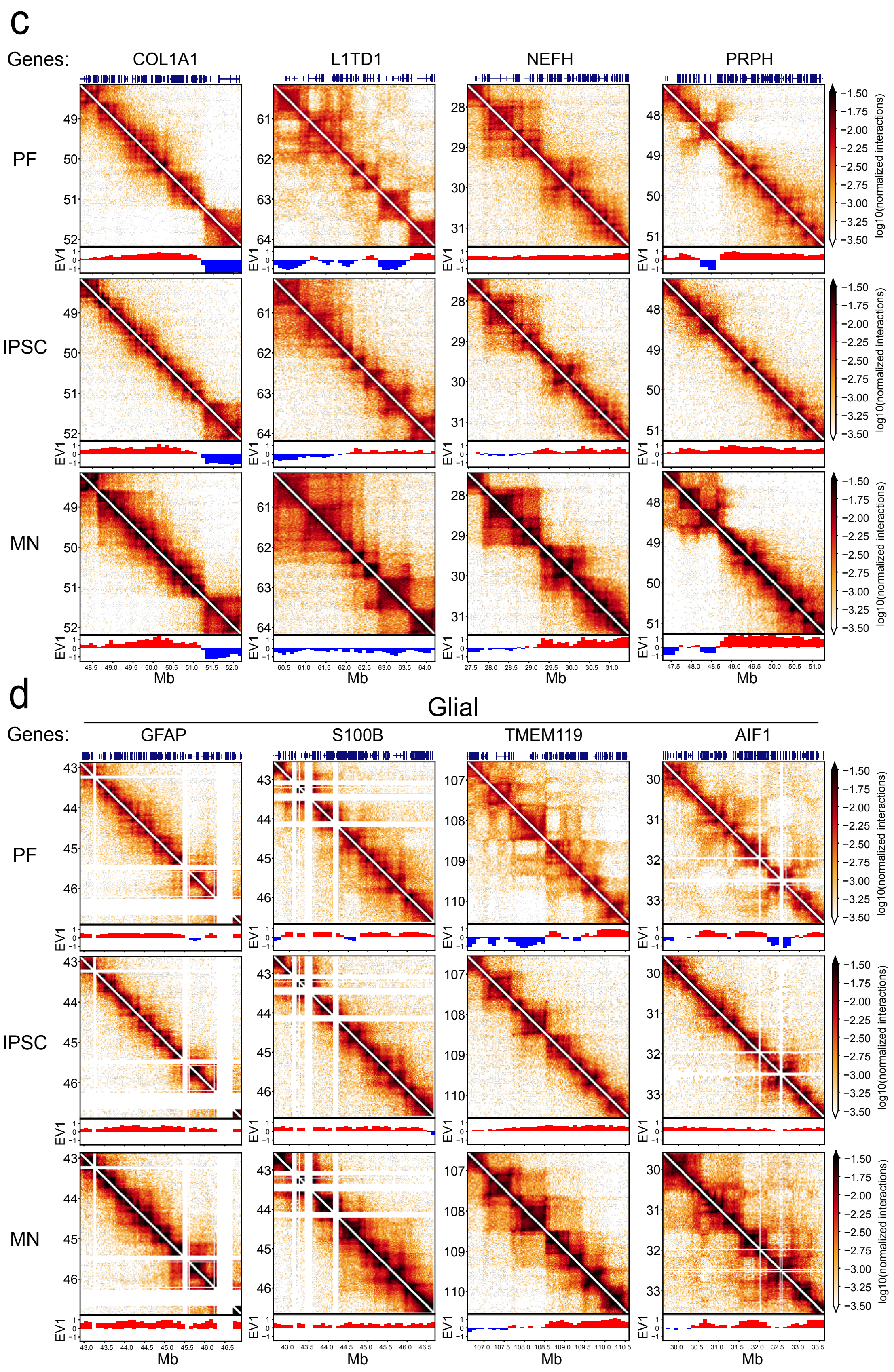

### Supplementary Figure S6

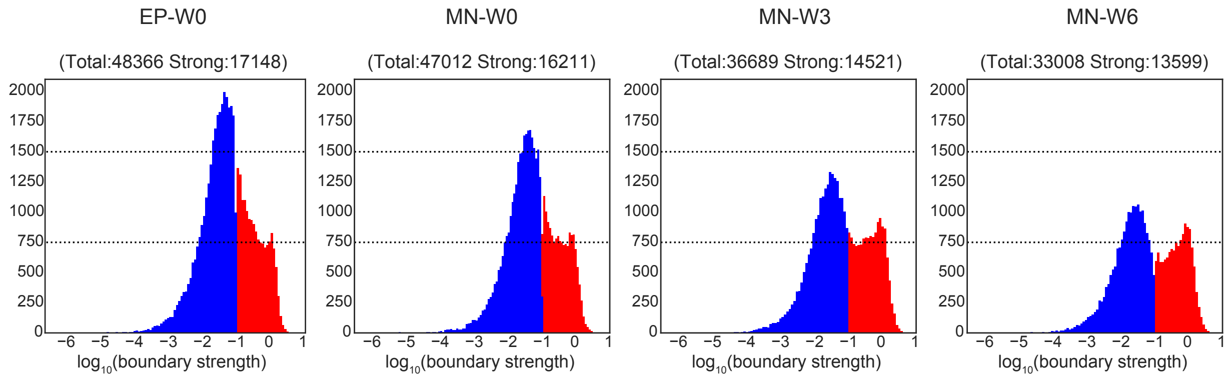

### Supplementary Figure S7AG

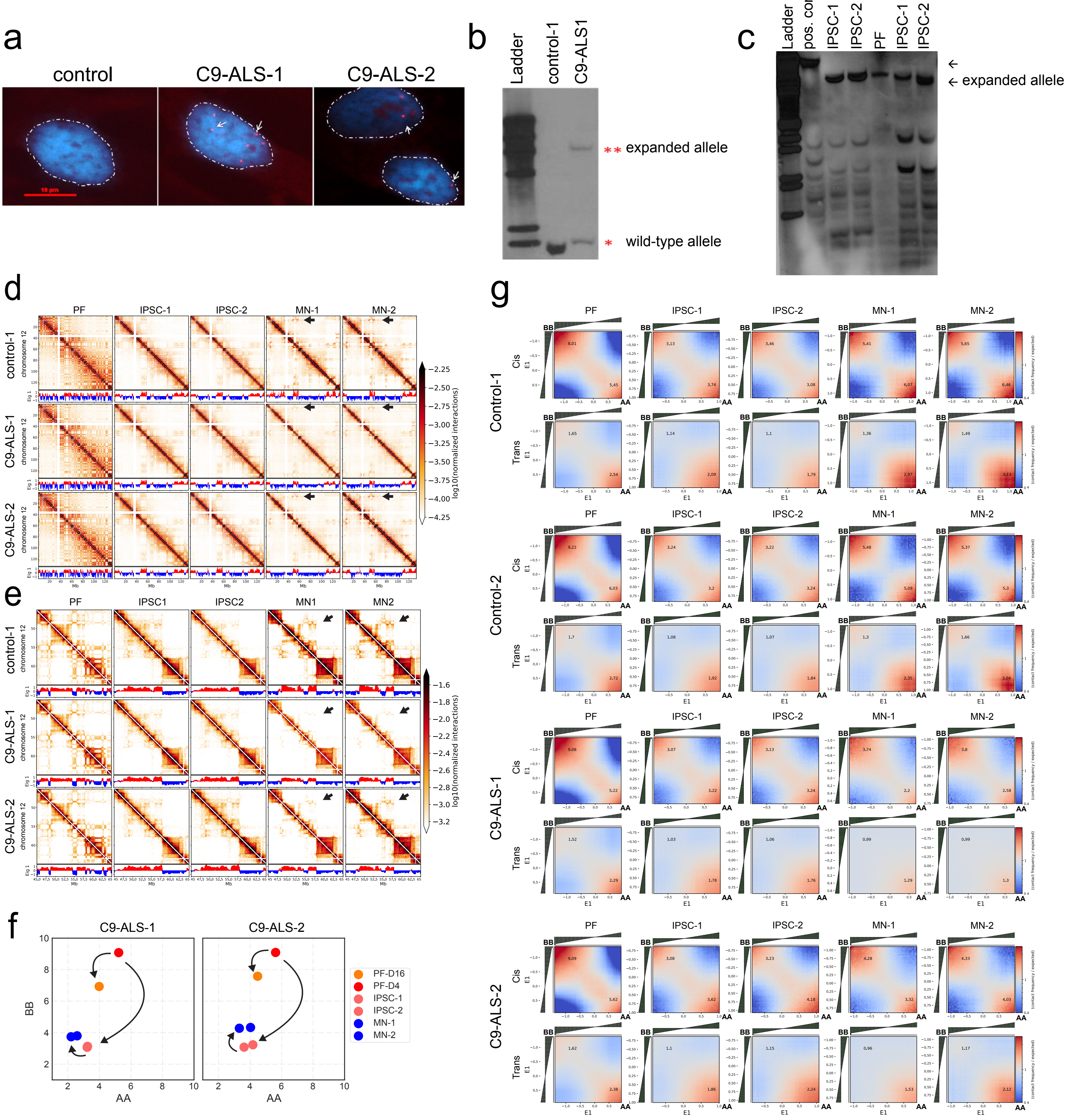

### Supplementary Figure S7HK

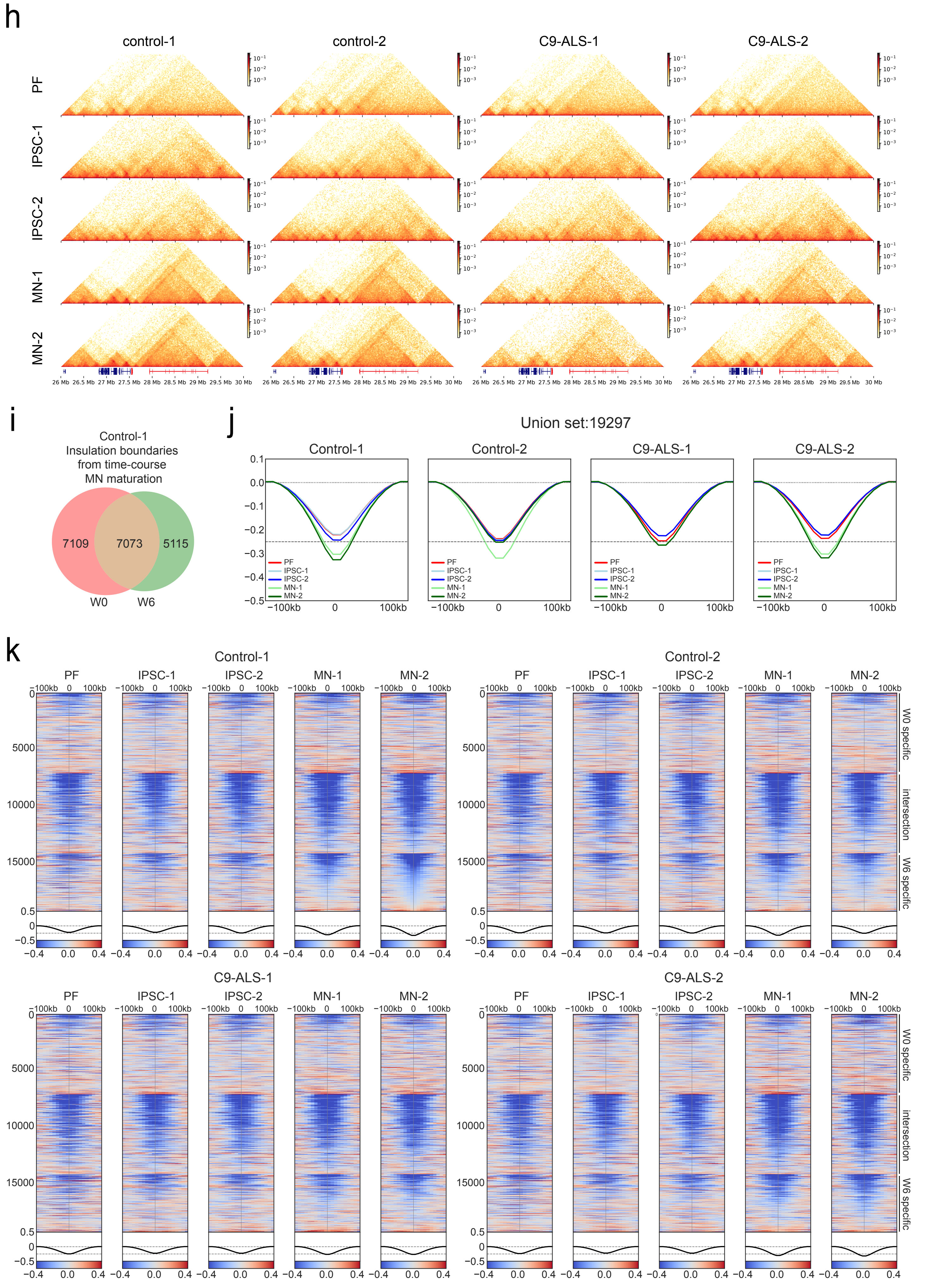

### Supplementary Figure S8AF

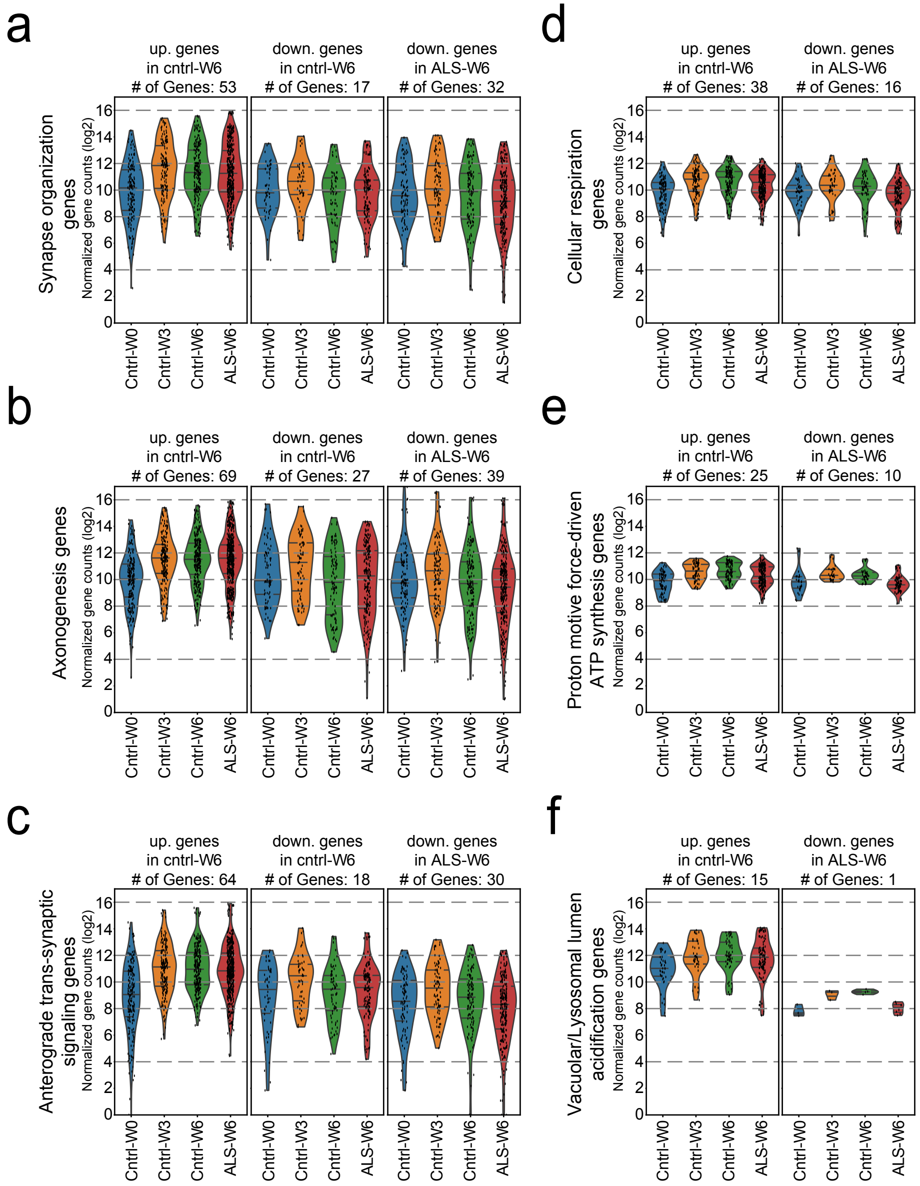

### Supplementary Figure S8g

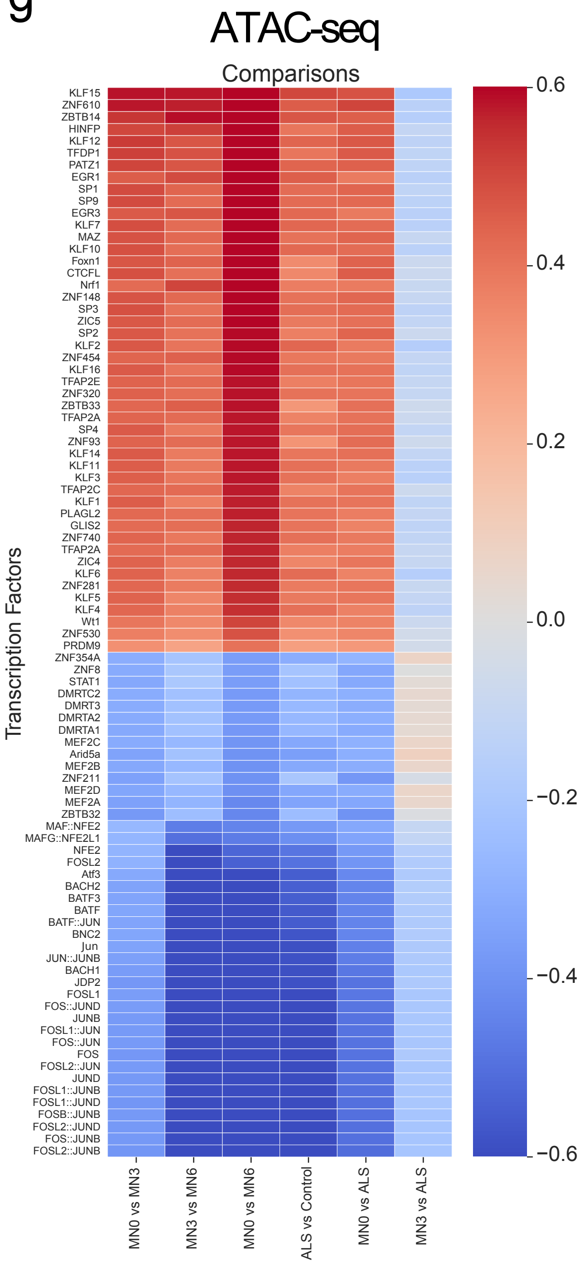
